# Sensory Tetanization to Induce LTP-Like Plasticity: A Review and Reassessment of the Approach

**DOI:** 10.1101/2022.03.06.483175

**Authors:** James W. Dias, Carolyn M. McClaskey, Jeffrey A. Rumschlag, Kelly C. Harris

**Affiliations:** Medical University of South Carolina

**Keywords:** auditory, high frequency stimulation, plasticity, sensory tetanization, visual, LTP

## Abstract

Great interest has been given to developing non-invasive approaches for studying cortical plasticity in humans. High frequency presentation of auditory and visual stimuli, or *sensory tetanization*, can induce long-term-potentiation-*like* (LTP-*like*) changes in cortical activity. However, contrasting effects across studies suggest that sensory tetanization may be unreliable. We review these contrasting effects, conduct our own study of auditory and visual tetanization, and perform meta-analyses to determine the average effect of sensory tetanization across studies. We measured auditory-evoked amplitude changes in a group of younger (18-29 years of age) and older (55-83 years of age) adults following tetanization to 1 kHz and 4 kHz tone bursts and following a slow-presentation control. We also measured visual-evoked amplitude changes following tetanization to horizontal and vertical sign gradients. Auditory and visual response amplitudes decreased following tetanization, consistent with some studies but contrasting with others finding amplitude increases (i.e., LTP-like changes). Older adults exhibited more modest auditory-evoked amplitude decreases, but visual-evoked amplitude decreases like those of younger adults. Changes in response amplitude were not specific to tetanized stimuli. Importantly, slow presentation of auditory tone-bursts produced response amplitude changes approximating those observed following tetanization in younger adults. Meta-analyses of visual and auditory tetanization studies found that the overall effect of sensory tetanization was not significant across studies or study sites. The results suggest that sensory tetanization may not produce reliable changes in cortical responses and more work is needed to determine the validity of sensory tetanization as a method for inducing human cortical plasticity in vivo.

## Introduction

Much attention has been given to the development of non-invasive techniques for inducing neuroplastic changes in the brain. Such techniques can be valuable for studying neural plasticity in humans and how neural plasticity is affected by environmental, neurochemical, clinical, and genetic factors. High frequency presentation of sensory stimuli, or *sensory tetanization* (Clapp, Kirk, et al., 2005; Teyler et al., 2005), has emerged as a popular technique for inducing changes in cortical activity that are often attributed to long-term potentiation (LTP) (for reviews, see Kirk et al., 2021; Sanders et al., 2018; Sumner, Spriggs, et al., 2020). While some have heralded sensory tetanization as a valuable non-invasive tool for the study of human cortical plasticity *in vivo* (Kirk et al., 2021; Sumner, Spriggs, et al., 2020), there are many inconsistent findings across studies that should be considered. Here, we provide background for the development and use of sensory tetanization as a technique for studying plasticity in the human brain. We then report our own study of auditory and visual tetanization in younger and older adults and compare our results to other studies. Finally, we report a meta-analysis to determine the extent to which sensory tetanization effects are consistent across studies and study sites.

## Approaches to Study Neural Plasticity

Long-term potentiation (LTP) describes the changes in synaptic structure that follow repeated neural stimulation that can facilitate efficient neural communication. First discovered in invertebrates (Kandel & Tauc, 1964, 1965) and later in vertebrate animals (Bliss & Lømo, 1973; Lømo, 1966; Lømo, 2003), LTP has emerged as a strong candidate neural mechanism for learning and memory (Bliss & Collingridge, 1993; Cooke & Bliss, 2006). Animal studies of LTP typically employ invasive techniques, including extracellular electrical stimulation of neural populations followed by local field recordings of population responses and post-mortem tissue analyses. Such invasive methods have limited the study of LTP in humans. As such, some of the most direct evidence for LTP in humans comes from studies demonstrating LTP in the resected cortical tissue of epileptic patients (e.g., Beck et al., 2000; Chen et al., 1996). Repetitive transcranial magnetic stimulation (rTMS) can non-invasively induce LTP-like changes in cortical activity in vivo, eliciting changes in cortical activity that reflect those observed in animal studies (for reviews, see Chung et al., 2016; e.g., Esser et al., 2006; Suppa et al., 2016; Wischnewski & Schutter, 2015). These changes in cortical activity are described as LTP-*like* because the changes in cortical activity after rTMS reflect those that accompany LTP changes in synaptic structure observed in animal studies, but synaptic structure is not (and cannot be) directly measured in vivo in humans.

Clapp, Kirk, et al. (2005) and Teyler et al. (2005) introduced another non-invasive method for inducing LTP-like plasticity by eliciting changes in cortical electrophysiological (EEG) activity after high-frequency presentation of auditory and visual stimuli. This *sensory tetanization* produced amplitude increases in the N1 and N1b components of auditory and visual evoked responses that they attributed to LTP in sensory cortex and described as LTP-like plasticity. Since the publication of these seminal studies, sensory tetanization has been used extensively to study plasticity in the human brain (for reviews, see Kirk et al., 2021; Sanders et al., 2018; Sumner, Spriggs, et al., 2020). Functional magnetic resonance imaging (fMRI) studies have reported changes in the BOLD response after tetanization that are localized in the sensory cortex (auditory or visual) coinciding with the modality of the test stimuli (Clapp, Zaehle, et al., 2005; Lahr et al., 2014; Wijtenburg et al., 2017; Zaehle et al., 2007). Magnetic resonance spectroscopy (MRS) studies have found that higher concentrations of excitatory neurotransmitters (i.e., glutamate) and lower concentrations of inhibitory neurotransmitters (i.e., γ-aminobutyric acid, GABA) are associated with larger event-related potential (ERP) amplitude increases following tetanization (Abuleil et al., 2019; Wijtenburg et al., 2017). Pharmaceuticals that can regulate these neurotransmitters can also modulate the effects of tetanization on ERP amplitudes (Forsyth et al., 2015; Sumner, McMillan, et al., 2020). Studies of aging find that older adults exhibit more modest changes in ERP amplitude after tetanization than younger adults (Abuleil et al., 2019; de Gobbi Porto et al., 2015; Spriggs et al., 2017). Studies of patients diagnosed with schizophrenia (Çavuş et al., 2012; D’Souza et al., 2018; Forsyth et al., 2017; Hamilton et al., 2020; Jahshan et al., 2017; Mears & Spencer, 2012; Valstad et al., 2021; but see Wynn et al., 2019), bipolar disorder (Elvsåshagen et al., 2012; Valstad et al., 2021; Zak et al., 2018), and depression (Normann et al., 2007; Sumner, McMillan, et al., 2020) have also found that these patient populations exhibit more modest changes in response amplitudes after tetanization compared to healthy controls, while adults with autism spectrum disorder can exhibit more pronounced response amplitude increases (Wilson et al., 2017). The broad interdisciplinary use of sensory tetanization to study plasticity and how plasticity changes across different experimental groups demonstrates the potential value of this in vivo approach. However, across these studies, a troubling degree of variability in the type and degree of change in neural activity elicited by sensory tetanization is evident, raising concerns regarding the reliability and validity of the approach to induce and study plasticity in the human brain.

## Inconsistent Changes in Response Amplitudes

The seminal studies introducing sensory tetanization as an in vivo approach to study plasticity reported an *increased* N1 and N1b amplitude in the sensory-evoked ERP after tetanization to auditory (Clapp, Kirk, et al., 2005) and visual (Teyler et al., 2005) stimuli, respectively. Subsequent studies have since replicated these results in young healthy adults, reporting increases in the N1 and N1b (and the C1, P1, P2) amplitudes of sensory-evoked ERPs after tetanization (Kleeva et al., 2022; Lei et al., 2017; Lengali et al., 2021; McNair et al., 2006; Moore et al., 2020; Moore & Loprinzi, 2021; Normann et al., 2007; Ross et al., 2008; Rygvold et al., 2020; Smallwood et al., 2015; Spriggs et al., 2017; Spriggs et al., 2018; Spriggs et al., 2019; Sumner et al., 2018; Wilson et al., 2017; Zak et al., 2018). However, despite the number of studies finding sensory ERP amplitudes increase after tetanization, several others have found no effect of tetanization on response amplitudes (D’Souza et al., 2018; Rygvold et al., 2020; Sumner et al., 2018) and several more have even found that response amplitudes *decrease* after tetanization (Kleeva et al., 2022; Klöppel et al., 2015; Rebreikina et al., 2021; Spriggs et al., 2018; Wynn et al., 2019). Such variability in response amplitude changes after sensory tetanization are not limited to the N1 or N1b components. C1 amplitude can increase (Rygvold et al., 2020), decrease (Forsyth et al., 2015; Lengali et al., 2021; Zak et al., 2018), or not change (Elvsåshagen et al., 2012) after tetanization. Similarly, P1 amplitude can increase (Elvsåshagen et al., 2012; Normann et al., 2007; Rygvold et al., 2020; Zak et al., 2018) or remain unchanged (Lengali et al., 2021; Teyler et al., 2005) after tetanization. P2 amplitude can also increase (Kleeva et al., 2022; Rygvold et al., 2020; Spriggs et al., 2017; Spriggs et al., 2018), decrease (Forsyth et al., 2015), or not change (Teyler et al., 2005) after tetanization. These contrasting effects of tetanization are also evident in fMRI studies, with three studies finding an increased BOLD response following tetanization (Clapp, Zaehle, et al., 2005; Wijtenburg et al., 2017; Zaehle et al., 2007) and one finding a decrease in BOLD response (Lahr et al., 2014). These variable findings are all reported in groups of younger healthy adults. For this review, we are not considering the variability reported in treatment groups or clinical samples.

These contrasting effects across studies raise concerns regarding the principal value of sensory tetanization as an in vivo method for studying neural plasticity. The initial finding that sensory tetanization increased ERP response amplitudes reflected the observations from animal studies that directly measured LTP, and so changes in ERP amplitudes after sensory tetanization were attributed to LTP in the human brain (Clapp, Kirk, et al., 2005; Teyler et al., 2005). However, findings across subsequent studies that employed methods similar to those initially reported by Clapp, Kirk, et al. (2005) and Teyler et al. (2005) suggest that sensory tetanization can result in response amplitude increases, decreases, or no change at all.

## Current Investigation

Though a large corpus of published studies has used sensory tetanization to study LTP-like plasticity in the human brain, contrasting findings across these studies suggest the approach may lack reliability and validity. Here we report a study of our own to determine whether sensory tetanization can elicit changes in the sensory-evoked potentials of younger and older adults. To provide a rigorous test of the specificity of changes in sensory-evoked potentials to tetanization, we incorporated a control condition absent of high-frequency stimulus presentation, instead exposing participants to a slow presentation of our test stimuli. We also incorporated different tetanizing and test stimuli to examine the degree to which changes in sensory-evoked potentials after tetanization were specific to the tetanized stimulus.

Following our study, our results and those reported in past studies were examined in meta-analyses to determine whether the average effect of visual and auditory tetanization on sensory-evoked ERPs was significant across studies. The studies included in our meta-analyses all used sensory tetanization to study LTP-like plasticity in the style of Clapp, Kirk, et al. (2005) and Teyler et al. (2005) and reported amplitude changes in the P1, N1, and P2 components of auditory-evoked ERPs and the C1, P1, N1, N1b, P2a, and P2 components of visual-evoked ERPs in groups of younger healthy adults (e.g., Clapp, Kirk, et al., 2005; Teyler et al., 2005). In the discussion, we also consider how studies not included in the meta-analyses support sensory tetanization as a reliable method for studying LTP-like plasticity in the human brain.

## Sensory Tetanization Study

### Methods

#### Participants

Our study sample consisted of 29 younger adults (18-29 years of age, M=22.52 years of age, SE=0.536, 20 women) and 33 older adults (55-83 years of age, M=67.61 years of age, SE=1.138, 22 women). Younger participants were graduate and undergraduate student volunteers from the Medical University of South Carolina and the College of Charleston in Charleston, SC. Older participants consisted of volunteers from an ongoing longitudinal study on presbycusis and volunteers from the greater Charleston, SC area. Several participants in both age groups had previous experience with hearing tasks, but all were unfamiliar with the auditory and visual tetanization tasks in the current study. All participants were native English speakers and had normal cognitive ability, having completed the Mini-Mental State Examination (Folstein et al., 1983) with no more than 3 errors. None of our participants were currently or recently taking psychopharmaceutical medications that affect cognition, including medications to treat depression, anxiety, AD/HD, or sleeping disorders. This study was approved by the Institutional Review Board of the Medical University of South Carolina and all participants provided written informed consent prior to participation.

Participants were limited to those with normal hearing or mild-to-moderate sensorineural hearing loss. Audiometric thresholds for conventional test frequencies are reported in Figure 1. The hearing thresholds of younger adults were within normal clinical limits (≤25 dB HL). The hearing thresholds of older adults were more variable, ranging from within normal clinical limits to mild-to-moderate high frequency hearing loss. Audiometric thresholds averaged across 500 to 4000 Hz were higher for older participants (M = 21.057 dB HL, SE = 2.186) than for younger participants (M = 1.807 dB HL, SE = 0.483), t(60) = −8.095, d = 9.343, p < 0.001. To identify any variance in the effects of auditory tetanization that may be accounted for by differences in audiometric thresholds, average thresholds were included in our statistical analyses.

**Figure 1.**
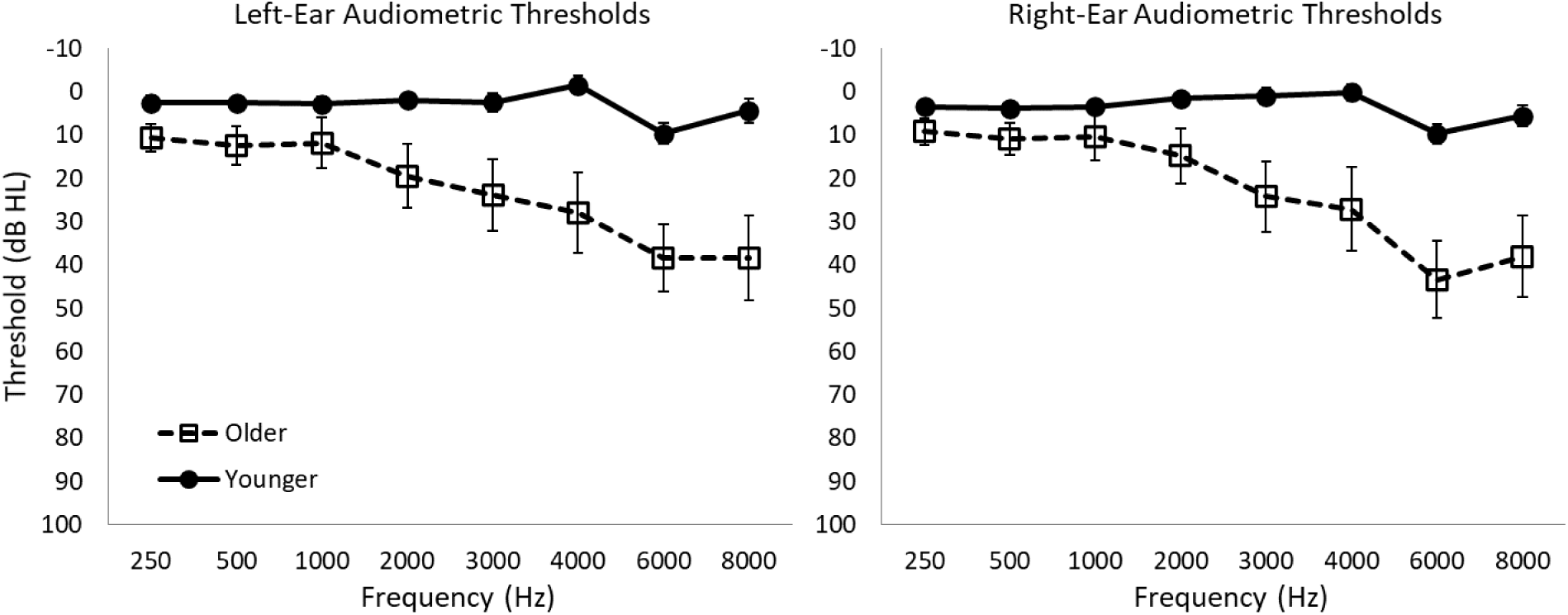
Audiometric thresholds for the left and right ears of younger and older adults. Error bars represent standard error (SE) of the mean.

Participants had a visual acuity ≤ 20/50 on the Snellen Eye Chart and a visual log contrast sensitivity ≥ 1.00 on the Pelli-Robson Contrast Sensitivity Chart. Younger adults had visual acuity between 20/10 and 20/25 (N = 27, M = 14.960, SE = 0.628) and visual log contrast sensitivity between 1.50 and 1.95 (N = 26, M = 1.685, SE = 0.017). Older adults had visual acuity between 20/13 and 20/50 (N = 29, M = 23.340, SE = 1.805) and visual log contrast sensitivity between 1.35 and 1.80 (N = 29, M = 1.557, SE = 0.023). There were significant age-group differences in visual acuity (t[54] = −4.260, d = 7.358, p < 0.001) and contrast sensitivity (t[53] = −4.381, d = 0.108, p < 0.001). Visual acuity and contrast sensitivity were included in our statistical analyses to determine whether these age-group differences in vision could account for variability in the effects of visual tetanization.

#### Auditory Tetanization

The auditory tetanization task was similar to procedures previously reported to elicit LTP-like changes in auditory ERP amplitudes (e.g., Clapp, Kirk, et al., 2005; Kleeva et al., 2022). The stimuli used to elicit auditory ERPs were 1 kHz and 4 kHz tone-bursts with a duration of 50 ms (including a 5-ms ramp on and a 5-ms ramp off) presented at 90 dB SPL. Baseline auditory ERPs were measured from two sessions of 220 presentations of 1 kHz and 4 kHz tone bursts. Each stimulus was randomly presented 110 times at a presentation frequency of ∼0.41-0.54 Hz – stimulus duration 50 ms and random ISI between 1800 and 2400 ms. Following baseline recording, participants were either tetanized to 1 kHz or 4 kHz tone bursts or repeated the baseline trial (i.e., the non-tetanized slow presentation of both 1 kHz and 4 kHz tone bursts). For the tetanization conditions, the tetanizing stimulus was presented for 4 minutes at a presentation frequency of 13.3 Hz – stimulus duration 50 ms and an ISI of 25 ms. Following tetanization or slow presentation, participants were given 5 minutes to rest. Post-tetanization and post-slow-presentation auditory ERPs were then measured following the same procedure used to record baseline auditory ERPs. Of our 29 younger participants, 20 completed 1 kHz tone-burst tetanization, 20 completed 4 kHz tone-burst tetanization, and 22 completed the slow presentation control. Of our 33 older participants, 28 completed 1 kHz tone-burst tetanization, 28 completed 4 kHz tone-burst tetanization, and 26 completed the slow presentation control. Tone-burst stimuli were delivered through ER-3C Etymotic insert earphones inserted into the left and right ears. Stimulus delivery and timing was controlled with TDT RPvds and a Tucker-Davis Technologies RZ6 auditory processing (Tucker-Davis Technologies, Alachua, FL).

#### Visual Tetanization

The visual tetanization task was similar to procedures previously reported to elicit LTP-like changes in visual ERP amplitudes (e.g., Smallwood et al., 2015; Spriggs et al., 2017). The stimuli used to elicit visual ERPs were two circular disks, each containing a black and white sine gradient with a size of 1000 x 1000 pixels and a spatial frequency of approximately 1.4 cycles per degree. These disks were oriented either horizontally or vertically, as illustrated in Figure 2. Baseline visual ERPs were measured from two sessions of 220 presentations of the horizontal and vertical stimuli. Each stimulus was randomly presented 110 times at a presentation frequency of ∼0.67-1 Hz – stimulus duration of 33 ms and a random ISI between 1000 and 1500 ms, accounting for the 60 Hz refresh rate of the computer screen. Following baseline recording, participants were tetanized to one of the two visual stimuli, either horizontal or vertical. The tetanizing stimulus was presented 2000 times at a presentation frequency of ∼8.6 Hz – stimulus duration of 33 ms and a random ISI between 67 and 100 ms, accounting for the 60 Hz refresh rate of the computer screen. Following tetanization, participants were given 5 minutes to rest and allow any afterimages of the tetanizing stimulus to dissipate. Post-tetanization visual ERPs were then measured following the same procedure used to record baseline visual ERPs. Visual tetanization was completed by 18 younger and 30 older participants. Visual stimuli were presented on a 27” Asus VN279 LCD computer monitor at a visual angle of approximately 14⁰. Stimulus delivery and timing was controlled with a combination of PsychoPy software (Peirce et al., 2019) and a Cedrus StimTracker Quad (Cedrus Corp., San Pedro, CA, United States) equipped with photodiodes for accurate timing of stimulus delivery.

**Figure 2.**
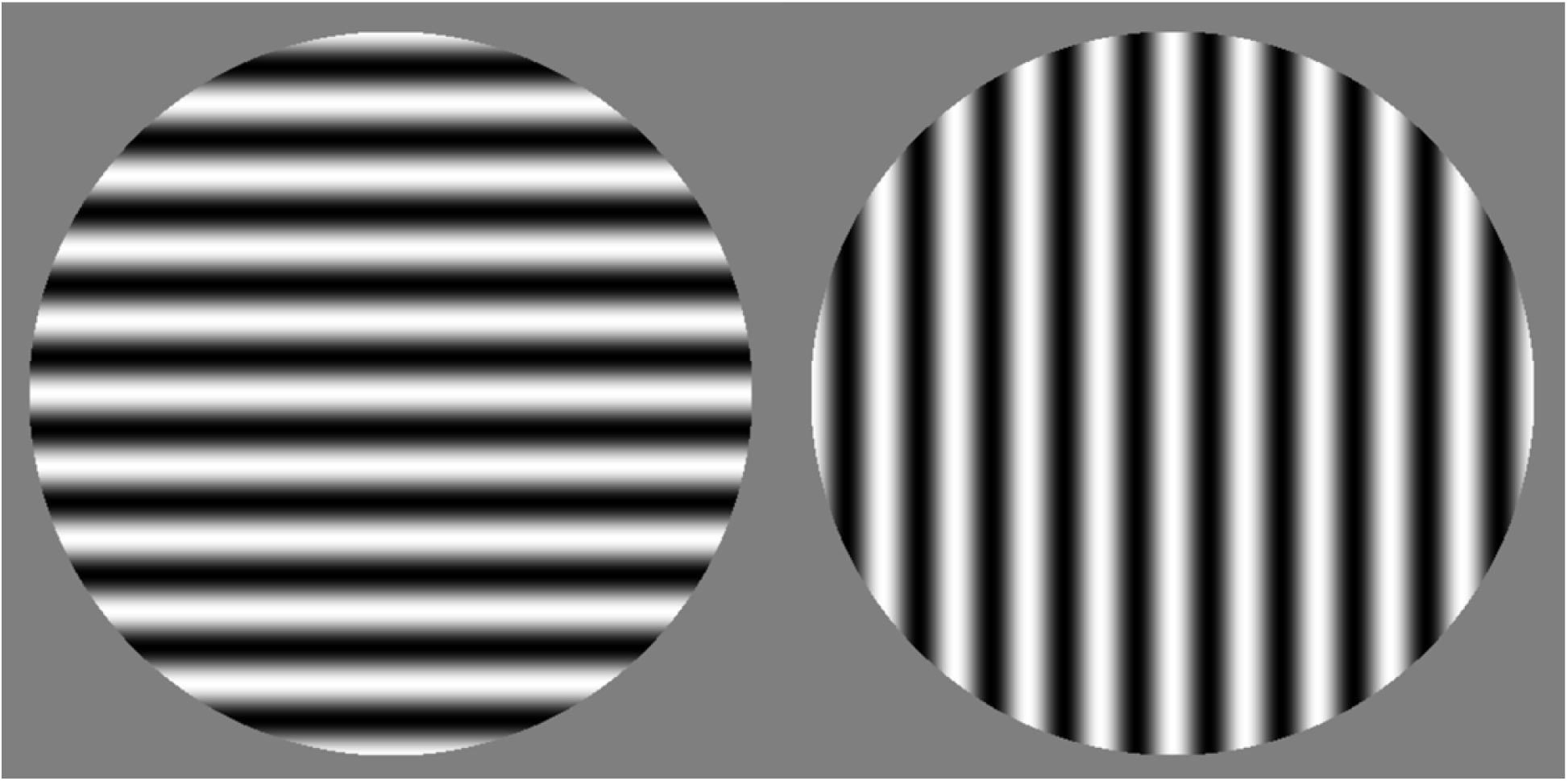
The horizontal (left) and vertical (right) sine gradients used in the visual tetanization task.

#### EEG Recording and ERP Analysis

Electroencephalography (EEG) was recorded from a 64-channel Neuroscan QuickCap (international 10-20 system) connected to a SynAmpsRT 64-Channel Amplifier. Bipolar electrodes placed above and below the left eye recorded vertical electro-oculogram activity and bipolar electrodes placed on the outer canthi of both eyes recorded horizontal electro-oculogram activity. Curry 8 software was used to record the EEG signal at a 10,000 Hz sampling rate while participants passively completed the auditory and visual tetanization tasks.

EEG data were then processed offline using a combination of EEGLab (Delorme & Makeig, 2004) and ERPLab (Lopez-Calderon & Luck, 2014). The recorded EEG data were down-sampled to 1000 Hz, bandpass filtered from 1-30 Hz, and re-referenced to the average of all electrodes. Auditory tetanization data were corrected for ocular artifacts by removing ocular source components following independent components analysis. Individual trials were then segmented into epochs around the stimulus onset (−100 ms – 500 ms) and then baseline corrected (−100 ms – 0 ms). Any epochs contaminated by peak-to-peak deflections in excess of 100 µV were rejected using an automatic artifact rejection algorithm. Visual tetanization data were also visually inspected to reject any epochs with amplitude deflections in the vertical and horizontal eye channels that could indicate blinks or eye-movements during the visual task.

For each participant, epoched data were averaged across trials to compute the average ERP waveform. ERPs were analyzed in a cluster of frontal-central electrodes for auditory data: F1, FZ, F2, FC1, FCZ, FC2, C1, CZ, C2 (e.g., Dias et al., 2018; Dias et al., 2021; Harris et al., 2012; Narne & Vanaja, 2008; Tremblay et al., 2001), and in a cluster of posterior-central electrodes for visual data: P7, TP7, CP5, P5, PO7, P8, TP8, CP6, P6, PO8 (McNair et al., 2006; Smallwood et al., 2015). ERPs were then averaged across channels before peak picking.

Post auditory stimulus onset, the latencies of the first prominent negative peak (N50), first prominent positive peak (P1), second prominent negative peak (N1), second prominent positive peak (P2), and third prominent negative peak (N2) were recorded. These latencies were used to compute temporal intervals for the automatic detection of the P1, N1, and P2 peak amplitudes and latencies in each of our channels of interest, individually, using custom MATLAB scripts. P1 was defined as the maximum positive peak between N50 (or from stimulus onset if no N50 was detected) and N1. N1 was defined as the maximum negative peak between P1 and P2. P2 was defined as the maximum positive peak between N1 and N2. P1, N1, and P2 peak amplitudes and latencies were then averaged across channels to yield a single metric for the P1 peak amplitude, P1 peak latency, N1 peak amplitude, N1 peak latency, P2 peak amplitude, and P2 peak latency for each participant. If a P1, N1, or P2 peak was not found in a channel between their respective latency windows defined from the average waveform across all of our channels of interest, then that channel was omitted from the computed average of each component characteristic (peak amplitude and peak latency). Representative examples of the P1, N1, and P2 components of the auditory-evoked response are provided in Figure 3.

**Figure 3.**
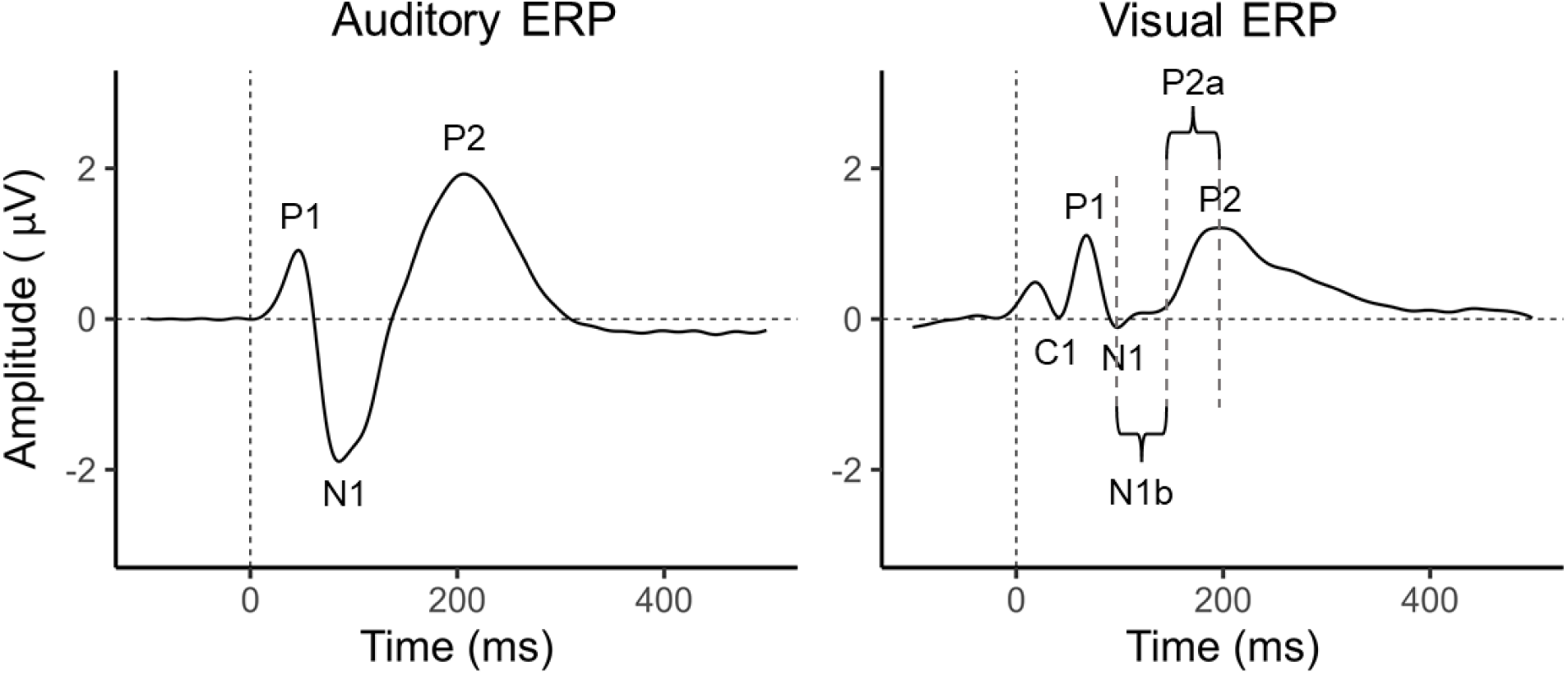
Representative examples of auditory and visual ERPs with components labeled.

Post visual stimulus onset, the latencies of the first prominent positive peak (P50), first prominent negative peak (C1), second prominent positive peak (P1), second prominent negative peak (N1), third prominent positive peak (P2), and third prominent negative peak (N2) were recorded. These latencies were used to compute temporal intervals for the automatic detection of the C1, P1, N1, and P2 peak amplitudes and latencies in each of our channels of interest, individually, using custom MATLAB scripts. C1 was defined as the maximum negative peak between P50 (or from stimulus onset if no P50 was detected) and P1. P1 was defined as the maximum positive peak between C1 and N1. N1 was defined as the maximum negative peak between P1 and P2. P2 was defined as the maximum positive peak between N1 and N2. C1, P1, N1, and P2 peak amplitudes and latencies were then averaged across channels to yield a single metric for the C1 peak amplitude, C1 peak latency, P1 peak amplitude, P1 peak latency, N1 peak amplitude, N1 peak latency, P2 peak amplitude, and P2 peak latency for each participant. If a C1, P1, N1, or P2 peak was not found in a channel between their respective latency windows defined from the average waveform across all of our channels of interest, then that channel was omitted from the computed average of each component characteristic (peak amplitude and peak latency). When a N1 and P2 amplitude were identified, the N1b and P2a amplitudes were computed. N1b amplitude is the average amplitude between N1 and the half-latency between N1 and P2. P2a amplitude is the average amplitude between P2 and the half latency between N1 and P2. N1b is the component of the visual-evoked response most often found to exhibit amplitude changes following visual tetanization (McNair et al., 2006; Moore et al., 2020; Ross et al., 2008; Smallwood et al., 2015; Spriggs et al., 2017; Teyler et al., 2005; Wynn et al., 2019). P2a, though not commonly investigated (Spriggs et al., 2017), may also exhibit changes following tetanization and so we included it in our analyses. Representative examples of the C1, P1, N1, N1b, P2a, and P2 components of the visual-evoked response are provided in Figure 3.

As with previous studies, this approach allowed for automatic peak picking across channels, reducing the risk of human error, while more accurately measuring components by not considering those channels that failed to exhibit an identifiable component (e.g., Anderer et al., 1996; Dias et al., 2018; Dias et al., 2021).

Individual and average auditory ERPs along with individual data for the P1, N1, and P2 components for slow presentation, 1 kHz tetanization, and 4 kHz tetanization are reported in Figure 4, Figure 5, and Figure 6, respectively. Individual and average visual ERPs along with individual data for the C1, P1, N1, N1b, P2a, and P2 components for horizontal and vertical tetanization are reported in Figure 7 and Figure 8, respectively. These figures illustrate the average ERP prior to tetanization (blue) and after tetanization (red) with the shaded area representing the standard error. Included in these figures are the pre-post tetanization waveforms for individual participants, represented as light red and light blue ribbons. Light blue shading of these ribbons highlights where a participant’s pre-tetanization waveform was larger than their post-tetanization waveform and light red (pink) shading highlights where a participant’s post-tetanization waveform was larger than their pre-tetanization waveform. Evident from these figures is a high degree of individual variability, with most participants exhibiting larger pre-tetanization amplitudes, but many others exhibiting larger post-tetanization amplitudes. Reported below these waveforms are box and dot plots representing the peak amplitudes of the ERP components measured from each waveform for each of our participants.

**Figure 4.**
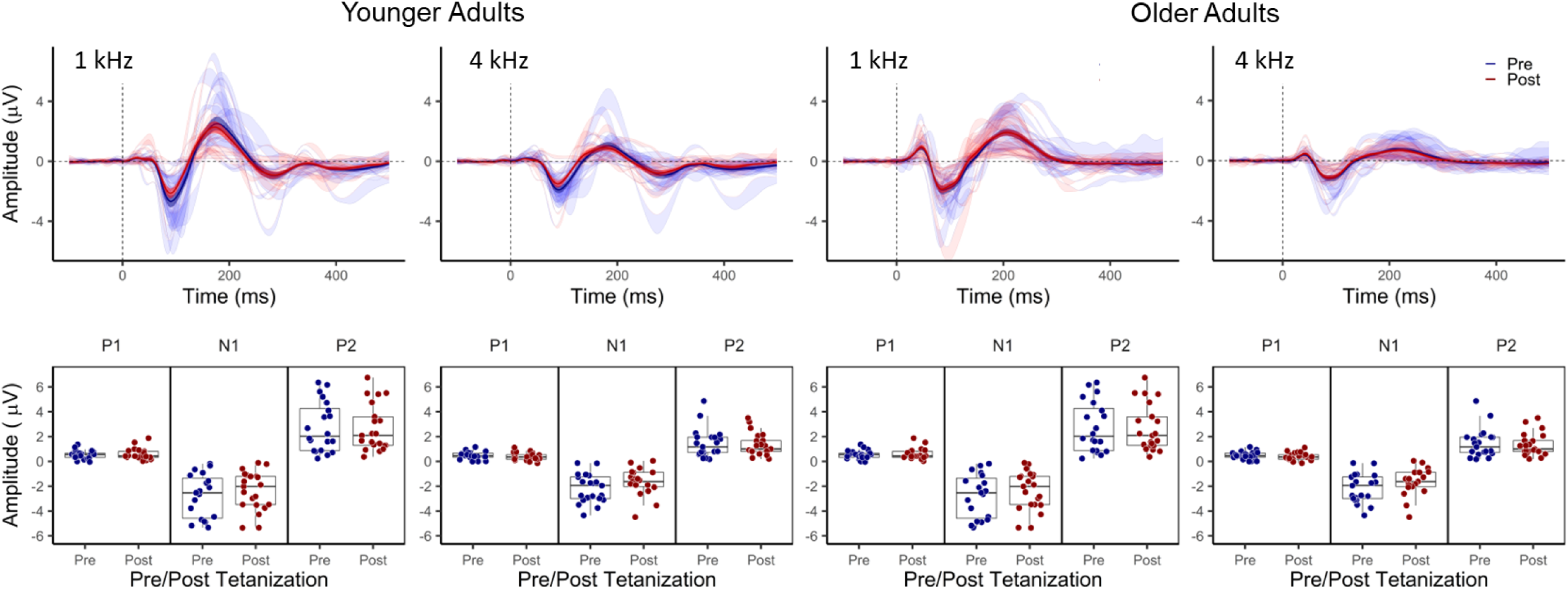
Effects of auditory tetanization by *slow presentation of 1 kHz and 4 kHz tone bursts*. Top panel shows average auditory-evoked ERPs generated by 1 kHz and 4 kHz tone bursts recorded in younger (left) and older (right) participants before and after *slow presentation of 1 kHz and 4 kHz tone bursts*. Solid blue = before tetanization, solid red = after tetanization. Shading around each solid line represents the standard error (SE) of the average waveform. The pre-post waveforms for individual participants are plotted as ribbons in the background of the average waveforms. Light blue shading of these ribbons highlights where a participant’s pre-tetanization waveform was larger than their post-tetanization waveform and light red (pink) shading highlights where a participant’s post-tetanization waveform was larger than their pre-tetanization waveform. Bottom panels below each waveform show the corresponding box and dot plots of individual response amplitudes for ERP components P1, N1, and P2.

**Figure 5.**
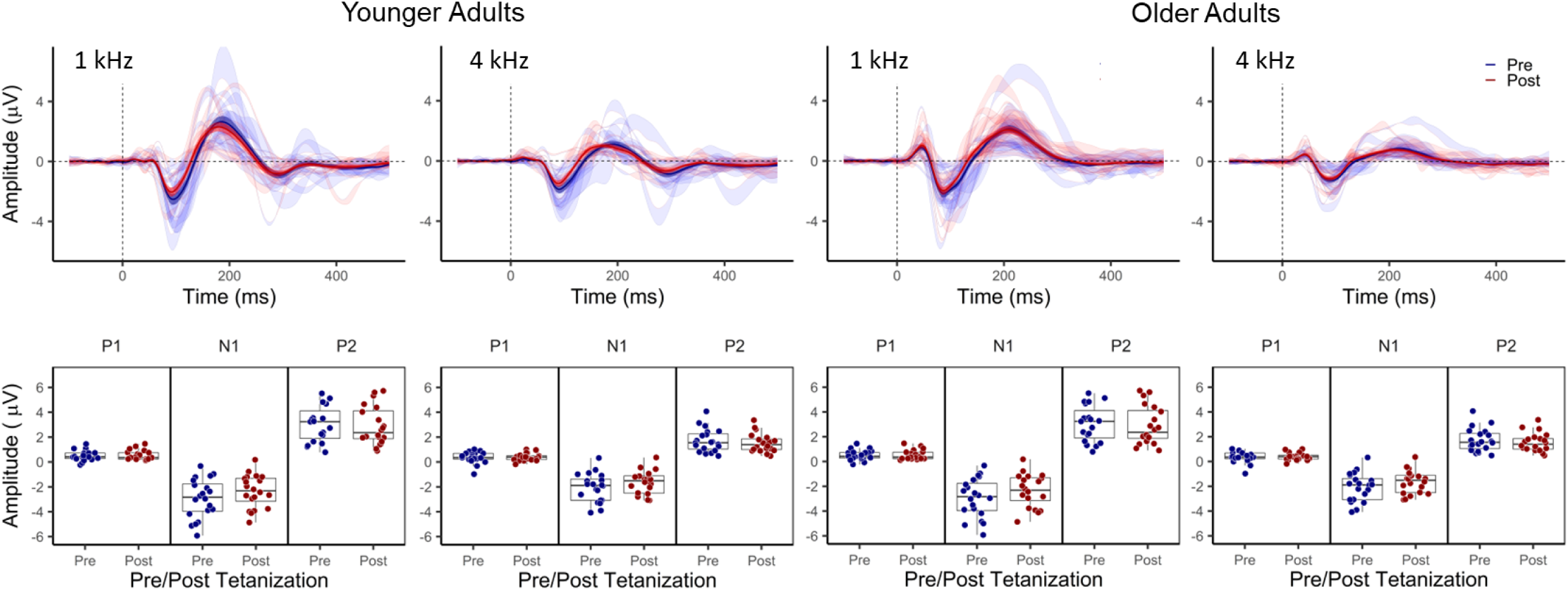
Effects of auditory tetanization by *1 kHz tone-bursts*. Top panel shows average auditory-evoked ERPs generated by 1 kHz and 4 kHz tone bursts recorded in younger (left) and older (right) participants before and after *1 kHz tone-burst tetanization*. Solid blue = before tetanization, solid red = after tetanization. Shading around each solid line represents the standard error (SE) of the average waveform. The pre-post waveforms for individual participants are plotted as ribbons in the background of the average waveforms. Light blue shading of these ribbons highlights where a participant’s pre-tetanization waveform was larger than their post-tetanization waveform and light red (pink) shading highlights where a participant’s post-tetanization waveform was larger than their pre-tetanization waveform. Bottom panels below each waveform show box and dot plots of individual response amplitudes for the corresponding ERP components P1, N1, and P2.

**Figure 6.**
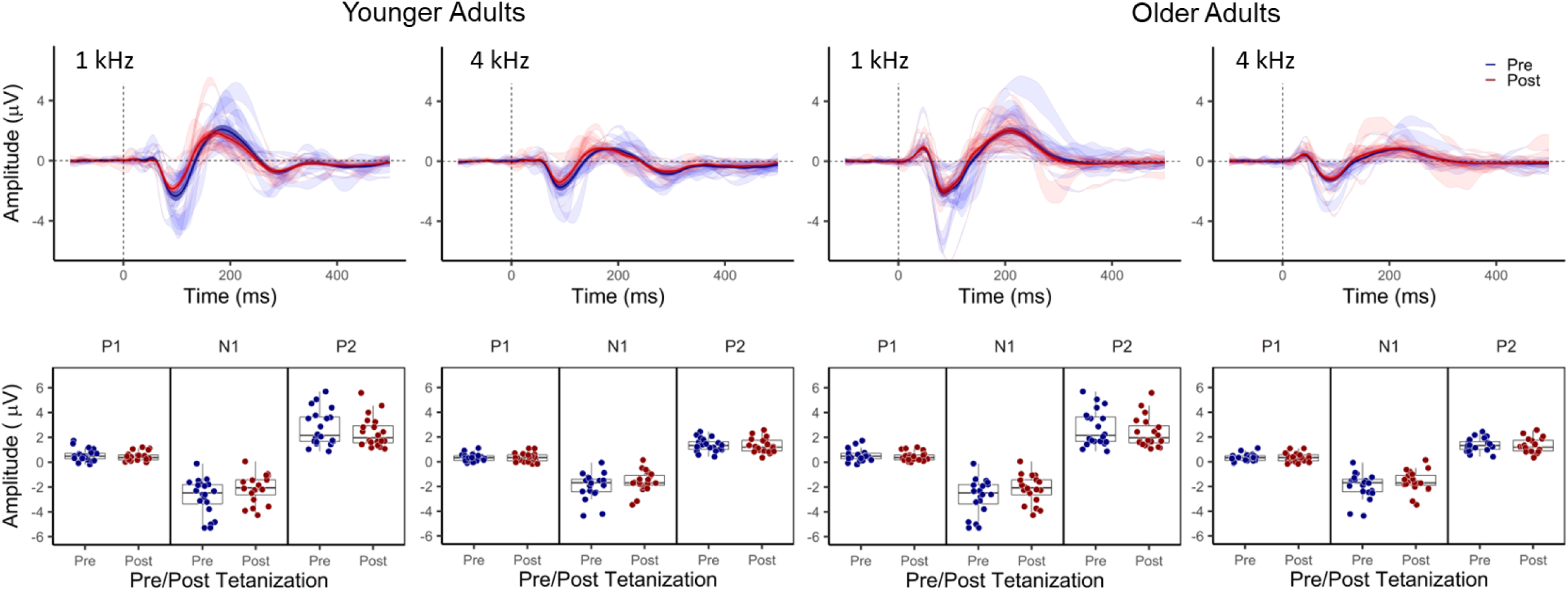
Effects of auditory tetanization by *4 kHz tone-*bursts. Top panel shows average auditory-evoked ERPs generated by 1 kHz and 4 kHz tone bursts recorded in younger (left) and older (right) participants before and after *4 kHz tone-burst tetanization*. Solid blue = before tetanization, solid red = after tetanization. Shading around each solid line represents the standard error (SE) of the average waveform. The pre-post waveforms for individual participants are plotted as ribbons in the background of the average waveforms. Light blue shading of these ribbons highlights where a participant’s pre-tetanization waveform was larger than their post-tetanization waveform and light red (pink) shading highlights where a participant’s post-tetanization waveform was larger than their pre-tetanization waveform. Bottom panels below each waveform show corresponding box and dot plots for individual response amplitudes of ERP components P1, N1, and P2.

**Figure 7.**
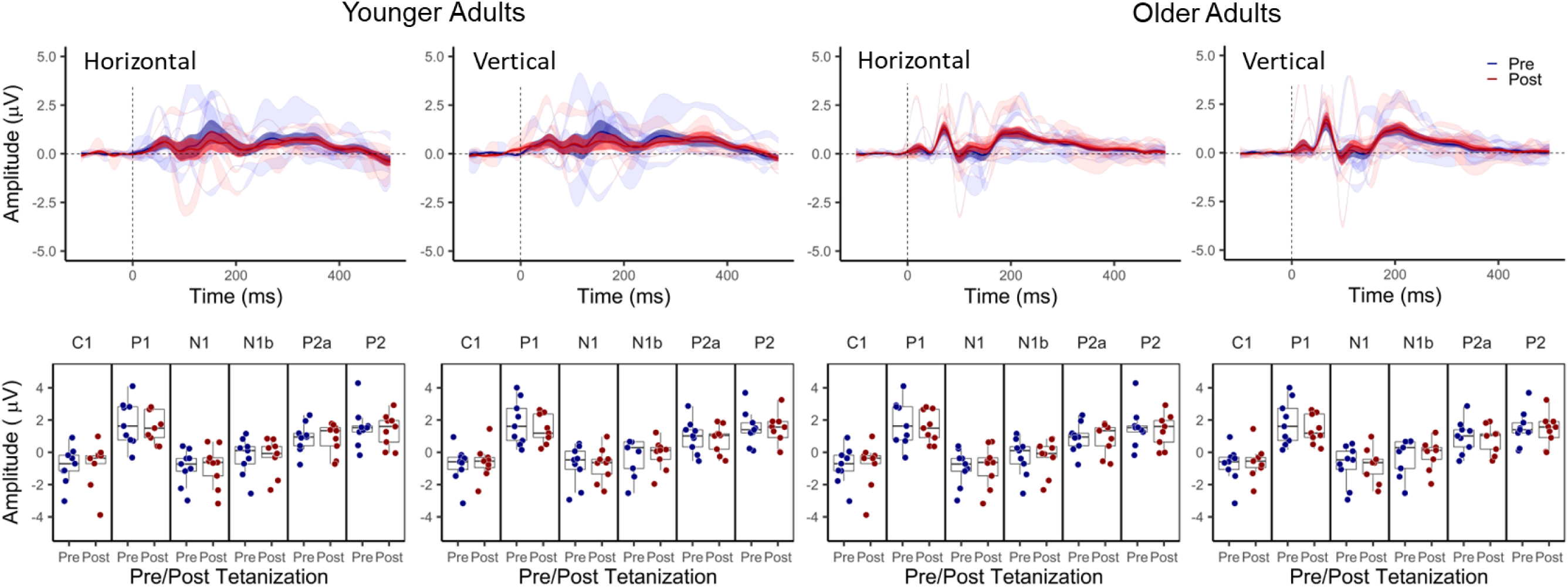
Effects of visual tetanization by a *horizontal* sign gradient. Top panel shows average visual-evoked ERPs generated by horizontal and vertical sine gradients recorded in younger (left) and older (right) participants before and after *horizontal tetanization*. Solid blue = before tetanization, solid red = after tetanization. Shading around each solid line represents the standard error (SE) of the average waveform. The pre-post waveforms for individual participants are plotted as ribbons in the background of the average waveforms. Light blue shading of these ribbons highlights where a participant’s pre-tetanization waveform was larger than their post-tetanization waveform and light red (pink) shading highlights where a participant’s post-tetanization waveform was larger than their pre-tetanization waveform. Bottom panels below each waveform show box and dot plots of individual response amplitudes for the corresponding ERP components C1, P1, N1, N1b, P2a, and P2.

**Figure 8.**
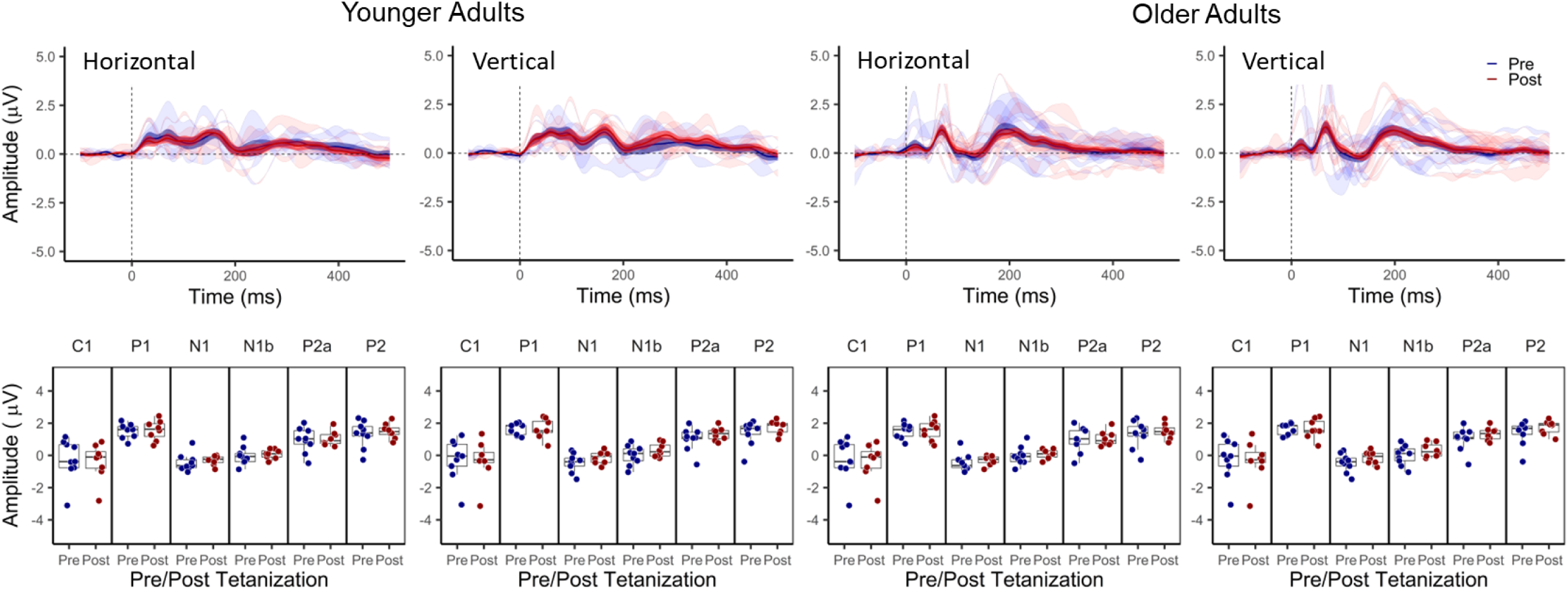
Effects of visual tetanization by a *vertical* sign gradient. Top panel shows average visual-evoked ERPs generated by horizontal and vertical sine gradients recorded in younger (left) and older (right) participants before and after *vertical tetanization*. Solid blue = before tetanization, solid red = after tetanization. Shading around each solid line represents the standard error (SE) of the average waveform. The pre-post waveforms for individual participants are plotted as ribbons in the background of the average waveforms. Light blue shading of these ribbons highlights where a participant’s pre-tetanization waveform was larger than their post-tetanization waveform and light red (pink) shading highlights where a participant’s post-tetanization waveform was larger than their pre-tetanization waveform. Bottom panels below each waveform show box and dot plots of individual response amplitudes for the corresponding ERP components C1, P1, N1, N1b, P2a, and P2.

#### Analytical Approach

Linear mixed effects regression (LMER) models were used to test for stimulus response amplitude changes following auditory and visual tetanization. LMER is a valuable non-parametric statistical approach that can test hypothesis-driven relationships while accounting for the variability in multi-level factors nested within participants (i.e., trial and stimulus).

LMER models tested for changes in auditory-evoked response amplitudes following 1 kHz tone-burst tetanization, 4 kHz tone-burst tetanization, and slow presentation of 1 kHz and 4 kHz tone bursts. P1, N1, and P2 peak amplitudes served as the outcome variables for separate LMER models. For each of these models, trial (pre-tetanization and post-tetanization), stimulus (1 kHz and 4kHz), and age group (younger and older) were included as fixed factors, including the interaction terms for trial and stimulus with age-group. Participant served as a random factor.

Similarly, LMER models tested for changes in visual-evoked response amplitudes following tetanization to horizontal and vertical sine-gradients. The fixed factors of trial and age group were included in the model, along with their interaction term, and participant served as a random factor. Tetanization condition (horizontal or vertical tetanization) and stimulus type (horizontal or vertical stimulus), along with their interactions with trial and age group, were then added to this model to determine whether effects of tetanization were stimulus specific.

### Results

#### Slow Presentation of 1 kHz and 4 kHz Tone Bursts

The LMER models for slow presentation of 1 kHz and 4 kHz stimuli are reported in Table 1. P1 peak amplitudes were larger for 1 kHz than for 4 kHz tone bursts and were larger for older than for younger participants, but P1 peak amplitudes were unaffected by slow stimulus presentation. N1 peak amplitudes were larger for 1 kHz than for 4 kHz tone bursts (more negative). N1 peak amplitudes were smaller (more positive) following slow stimulus presentation and this effect of slow stimulus presentation interacted with age group. Post-hoc comparisons found that N1 peak amplitudes decreased following slow presentation for younger adults, but not for older adults. P2 peak amplitudes were larger for 1 kHz tone bursts than for 4 kHz tone bursts, but were otherwise unaffected by slow stimulus presentation or age-group. Adding audiometric thresholds to these models did not change the pattern of significant relationships or improve model fit (p>0.1).

**Table 1.** Slow Presentation Control Linear Mixed Effects Models of N1 Peak Ampitude Predicted by Stimulus, Trial, and Age Group

| | *B* | *SE* | $\beta$ | *SE* | *t* | *p* | |
| --- | --- | --- | --- | --- | --- | --- | --- |
| P1 Across Age-Groups |  |  |  |  |  |  |  |
| Stimulus 1 kHz / 4 kHz | -0.325 | 0.049 | -0.534 | 0.080 | -6.688 | <0.001 | *** |
| Pre/Post | 0.042 | 0.062 | 0.069 | 0.102 | 0.672 | 0.502 |  |
| Age Group Y / O | 0.312 | 0.140 | 0.512 | 0.230 | 2.225 | 0.030 | * |
| Pre/Post x Stimulus | -0.077 | 0.069 | -0.127 | 0.113 | -1.120 | 0.263 |  |
| Pre/Post x AgeGroup | 0.045 | 0.070 | 0.074 | 0.114 | 0.643 | 0.521 |  |
| N1 Across Age-Groups |  |  |  |  |  |  |  |
| Stimulus 1 kHz / 4 kHz | 0.926 | 0.091 | 0.631 | 0.062 | 10.145 | <0.001 | *** |
| Pre/Post | 0.703 | 0.117 | 0.479 | 0.080 | 5.997 | <0.001 | *** |
| Age Group Y / O | 0.723 | 0.370 | 0.493 | 0.252 | 1.954 | 0.056 |  |
| Pre/Post x Stimulus | -0.139 | 0.129 | -0.095 | 0.088 | -1.078 | 0.282 |  |
| Pre/Post x AgeGroup | -0.515 | 0.131 | -0.351 | 0.089 | -3.932 | <0.001 | *** |
| N1 Younger |  |  |  |  |  |  |  |
| Stimulus 1 kHz / 4 kHz | 0.905 | 0.162 | 0.572 | 0.102 | 5.601 | <0.001 | *** |
| Pre/Post | 0.735 | 0.164 | 0.464 | 0.103 | 4.489 | <0.001 | *** |
| Pre/Post x Stimulus | -0.204 | 0.230 | -0.129 | 0.145 | -0.888 | 0.376 |  |
| N1 Older |  |  |  |  |  |  |  |
| Stimulus 1 kHz / 4 kHz | 0.942 | 0.101 | 0.711 | 0.076 | 9.360 | <0.001 | *** |
| Pre/Post | 0.162 | 0.101 | 0.122 | 0.076 | 1.607 | 0.110 |  |
| Pre/Post x Stimulus | -0.087 | 0.142 | -0.066 | 0.107 | -0.614 | 0.540 |  |
| P2 Across Age-Groups |  |  |  |  |  |  |  |
| Stimulus 1 kHz / 4 kHz | -1.291 | 0.103 | -0.867 | 0.069 | -12.546 | <0.001 | *** |
| Pre/Post | -0.195 | 0.132 | -0.131 | 0.089 | -1.478 | 0.140 |  |
| Age Group Y / O | -0.582 | 0.343 | -0.391 | 0.230 | -1.697 | 0.096 |  |
| Pre/Post x Stimulus | 0.021 | 0.146 | 0.014 | 0.098 | 0.147 | 0.883 |  |
| Pre/Post x AgeGroup | 0.143 | 0.148 | 0.096 | 0.099 | 0.966 | 0.335 |  |

Notes: \* $p < 0.05$ , \*\* $p < 0.01$ , \*\*\* $p < 0.001$ . All significant values remained significant after correcting for inflated family-wise error using the Benjamini-Hochberg Procedure.

#### 1 kHz Tone-Burst Tetanization

The LMER models for 1 kHz tone-burst tetanization are reported in Table 2. P1 peak amplitudes were larger for 1 kHz than for 4 kHz tone bursts and were larger for older participants, but P1 peak amplitudes were unaffected by 1 kHz tone-burst tetanization. N1 peak amplitudes were larger for 1 kHz than for 4 kHz tone bursts (more negative). N1 peak amplitudes were smaller (more positive) following 1 kHz tone-burst tetanization and this effect of tetanization interacted with age group. Post-hoc comparisons found that N1 peak amplitudes decreased following 1 kHz tone-burst tetanization for younger and older adults, but the effect of tetanization was greater for younger adults. P2 peak amplitudes were larger for 1 kHz tone bursts than for 4 kHz tone bursts and were larger for younger adults than for older adults. P2 peak amplitudes were also smaller following tetanization to 1 kHz tone bursts, but this effect of tetanization did not interact with stimulus or age group. Adding audiometric thresholds to these models did not change the pattern of significant relationships or improve model fit (p>0.1).

**Table 2.** 1 kHz Tetanization Linear Mixed Effects Models of N1 Peak Ampitude Predicted by Stimulus, Trial, and Age Group

| | *B* | *SE* | $\beta$ | *SE* | *t* | *p* | |
| --- | --- | --- | --- | --- | --- | --- | --- |
| P1 Across Age-Groups |  |  |  |  |  |  |  |
| Stimulus 1 kHz / 4 kHz | -0.371 | 0.055 | -0.544 | 0.080 | -6.792 | <0.001 | *** |
| Pre/Post | 0.036 | 0.071 | 0.053 | 0.105 | 0.509 | 0.611 |  |
| Age Group Y / O | 0.416 | 0.155 | 0.611 | 0.228 | 2.680 | 0.010 | ** |
| Pre/Post x Stimulus | -0.022 | 0.077 | -0.032 | 0.113 | -0.279 | 0.781 |  |
| Pre/Post x AgeGroup | 0.065 | 0.078 | 0.096 | 0.115 | 0.834 | 0.405 |  |
| N1 Across Age-Groups |  |  |  |  |  |  |  |
| Stimulus 1 kHz / 4 kHz | 0.998 | 0.080 | 0.752 | 0.060 | 12.541 | <0.001 | *** |
| Pre/Post | 0.604 | 0.104 | 0.456 | 0.078 | 5.825 | <0.001 | *** |
| Age Group Y / O | 0.531 | 0.335 | 0.400 | 0.253 | 1.583 | 0.120 |  |
| Pre/Post x Stimulus | -0.190 | 0.113 | -0.144 | 0.085 | -1.692 | 0.092 |  |
| Pre/Post x AgeGroup | -0.316 | 0.114 | -0.238 | 0.086 | -2.770 | 0.006 | ** |
| N1 Younger |  |  |  |  |  |  |  |
| Stimulus 1 kHz / 4 kHz | 0.873 | 0.120 | 0.644 | 0.088 | 7.299 | <0.001 | *** |
| Pre/Post | 0.600 | 0.120 | 0.442 | 0.088 | 5.010 | <0.001 | *** |
| Pre/Post x Stimulus | -0.181 | 0.169 | -0.133 | 0.125 | -1.069 | 0.287 |  |
| N1 Older |  |  |  |  |  |  |  |
| Stimulus 1 kHz / 4 kHz | 1.086 | 0.106 | 0.846 | 0.082 | 10.256 | <0.001 | *** |
| Pre/Post | 0.291 | 0.106 | 0.227 | 0.082 | 2.751 | 0.007 | ** |
| Pre/Post x Stimulus | -0.197 | 0.150 | -0.153 | 0.117 | -1.315 | 0.190 |  |
| P2 Across Age-Groups |  |  |  |  |  |  |  |
| Stimulus 1 kHz / 4 kHz | -1.373 | 0.099 | -0.976 | 0.070 | -13.915 | <0.001 | *** |
| Pre/Post | -0.312 | 0.129 | -0.222 | 0.091 | -2.424 | 0.016 | * |
| Age Group Y / O | -0.751 | 0.309 | -0.534 | 0.220 | -2.432 | 0.019 | * |
| Pre/Post x Stimulus | 0.052 | 0.140 | 0.037 | 0.099 | 0.370 | 0.711 |  |
| Pre/Post x AgeGroup | 0.242 | 0.142 | 0.172 | 0.101 | 1.710 | 0.088 |  |

Notes: \* $p < 0.05$ , \*\* $p < 0.01$ , \*\*\* $p < 0.001$ . All significant values remained significant after correcting for inflated family-wise error using the Benjamini-Hochberg Procedure.

#### 4 kHz Tone-Burst Tetanization

The LMER models for 4 kHz tone-burst tetanization are reported in Table 3. P1 peak amplitudes were larger for 1 kHz than for 4 kHz tone bursts and were larger for older participants, but P1 peak amplitudes were unaffected by 4 kHz tone-burst tetanization. N1 peak amplitudes were larger for 1 kHz than for 4 kHz tone bursts (more negative). N1 peak amplitudes were smaller (more positive) following 4 kHz tone-burst tetanization and this effect of tetanization interacted with age group. Post-hoc comparisons found that N1 peak amplitudes decreased following 4 kHz tone-burst tetanization for younger and older adults, but the effect of tetanization was greater for younger adults. P2 peak amplitudes were larger for 1 kHz tone bursts than for 4 kHz tone bursts and were larger for younger adults than for older adults. P2 peak amplitudes were also smaller following tetanization to 4 kHz tone bursts, but this effect of tetanization did not interact with stimulus or age group. Adding audiometric thresholds to these models did not change the pattern of significant relationships or improve model fit (p>0.1).

**Table 3.** 4 kHz Tetanization Linear Mixed Effects Models of N1 Peak Ampitude Predicted by Stimulus, Trial, and Age Group

| | *B* | *SE* | $\beta$ | *SE* | *t* | *p* | |
| --- | --- | --- | --- | --- | --- | --- | --- |
| P1 Across Age-Groups |  |  |  |  |  |  |  |
| Stimulus 1 kHz / 4 kHz | -0.398 | 0.045 | -0.648 | 0.073 | -8.856 | <0.001 | *** |
| Pre/Post | -0.063 | 0.059 | -0.103 | 0.096 | -1.073 | 0.284 |  |
| Age Group Y / O | 0.366 | 0.142 | 0.595 | 0.231 | 2.573 | 0.013 | * |
| Pre/Post x Stimulus | 0.039 | 0.064 | 0.063 | 0.103 | 0.609 | 0.543 |  |
| Pre/Post x AgeGroup | 0.100 | 0.065 | 0.163 | 0.105 | 1.549 | 0.122 |  |
| N1 Across Age-Groups |  |  |  |  |  |  |  |
| Stimulus 1 kHz / 4 kHz | 0.964 | 0.077 | 0.737 | 0.059 | 12.548 | <0.001 | *** |
| Pre/Post | 0.590 | 0.101 | 0.451 | 0.077 | 5.841 | <0.001 | *** |
| Age Group Y / O | 0.376 | 0.335 | 0.288 | 0.257 | 1.122 | 0.267 |  |
| Pre/Post x Stimulus | -0.206 | 0.109 | -0.158 | 0.083 | -1.901 | 0.058 |  |
| Pre/Post x AgeGroup | -0.320 | 0.111 | -0.245 | 0.085 | -2.891 | 0.004 | ** |
| N1 Younger |  |  |  |  |  |  |  |
| Stimulus 1 kHz / 4 kHz | 0.812 | 0.122 | 0.659 | 0.099 | 6.660 | <0.001 | *** |
| Pre/Post | 0.656 | 0.122 | 0.533 | 0.099 | 5.370 | <0.001 | *** |
| Pre/Post x Stimulus | -0.338 | 0.172 | -0.275 | 0.140 | -1.961 | 0.052 |  |
| N1 Older |  |  |  |  |  |  |  |
| Stimulus 1 kHz / 4 kHz | 1.069 | 0.096 | 0.790 | 0.071 | 11.123 | <0.001 | *** |
| Pre/Post | 0.224 | 0.096 | 0.165 | 0.071 | 2.329 | 0.021 | * |
| Pre/Post x Stimulus | -0.115 | 0.136 | -0.085 | 0.100 | -0.845 | 0.399 |  |
| P2 Across Age-Groups |  |  |  |  |  |  |  |
| Stimulus 1 kHz / 4 kHz | -1.284 | 0.087 | -1.078 | 0.073 | -14.727 | <0.001 | *** |
| Pre/Post | -0.301 | 0.115 | -0.253 | 0.096 | -2.626 | 0.009 | ** |
| Age Group Y / O | -0.355 | 0.263 | -0.298 | 0.221 | -1.348 | 0.184 |  |
| Pre/Post x Stimulus | 0.185 | 0.123 | 0.155 | 0.103 | 1.501 | 0.134 |  |
| Pre/Post x AgeGroup | 0.239 | 0.126 | 0.201 | 0.105 | 1.906 | 0.058 |  |

Notes: \* $p < 0.05$ , \*\* $p < 0.01$ , \*\*\* $p < 0.001$ . All significant values remained significant after correcting for inflated family-wise error using the Benjamini-Hochberg Procedure.

#### Comparing Results Across Auditory Conditions

Across our experimental manipulations (slow presentation, 1 kHz tone-burst tetanization, and 4 kHz tone-burst tetanization), only N1 and P2 peak amplitudes changed significantly following tetanization, each decreasing in amplitude. Importantly, these effects of tetanization did not interact with stimulus, indicating that the effects of tetanization are not specific to the tetanizing stimulus. Additionally, younger adults demonstrated similar changes in N1 and P2 peak amplitudes following slow presentation, suggesting that changes in response amplitudes did not depend on a high stimulus-presentation rate.

To determine whether the effects of trial on N1 peak amplitude differed across our models, we performed Z-tests of the trial-coefficient differences between the auditory conditions for each age-group. Z scores for the coefficient differences were calculated as:

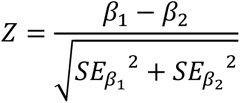

where *β_1_* and *β_2_* are the standardized coefficients from two different models that were compared and *SE*_β1_ and *SE*_β2_ are the standard errors for the respective standardized coefficients (Clogg et al., 1995; Cohen et al., 2003; Dias et al., 2021; McClaskey et al., 2018; Paternoster et al., 1998).

For younger adults, slow presentation of the 1 kHz and 4 kHz stimuli produced a decrease in N1 peak amplitude that was not significantly different from the decreases in N1 peak amplitude following 1 kHz tone-burst tetanization (Z = 0.162, p = 0.435) and 4 kHz tone-burst tetanization (Z = −0.483, p = 0.315). There was also no difference in the decrease in N1 peak amplitude following 1 kHz tone-burst tetanization and 4 kHz tone-burst tetanization (Z = −0.687, p = 0.246).

Interestingly, even though the LMER models reported above suggest that tetanization is needed to elicit decreases in N1 peak amplitude in older adults, slow presentation of the 1 kHz and 4 kHz stimuli produced a decrease in N1 peak amplitude that was not significantly different from the decreases in N1 peak amplitude following 1 kHz tone-burst tetanization (Z = −0.936, p = 0.175) and 4 kHz tone-burst tetanization (Z = −0.418, p = 0.338). This suggests that, for older adults, although the decrease in N1 peak amplitude following slow presentation was not significant, this decrease was also not significantly different from the significant decreases in N1 peak amplitude following 1 kHz and 4 kHz tetanization. Additionally, N1 peak amplitude decreases following 1 kHz tone-burst tetanization and 4 kHz tone-burst tetanization did not significantly differ (Z = 0.565, p = 0.286).

We performed additional Z-tests to determine whether the effects of trial on P2 peak amplitude across age-groups differed between the slow-presentation and 1 kHz and 4 kHz tetanization conditions. Slow presentation of the 1 kHz and 4 kHz stimuli produced a decrease in P2 peak amplitude that was not significantly different from the decreases in P2 peak amplitude following 1 kHz tone-burst tetanization (Z = 0.711, p = 0.238) and 4 kHz tone-burst tetanization (Z = 0.928, p = 0.177). There was also no difference in the decrease in P2 peak amplitude following 1 kHz tone-burst tetanization and 4 kHz tone-burst tetanization (Z = 0.232, p = 0.408).

The results suggest, overall, that slowly presenting tone-burst stimuli produced changes in N1 and P2 peak amplitude that were similar to those produced by tetanization for both younger and older adults.

#### Visual Tetanization

Two younger and two older participants were excluded from analyses for having response amplitudes that were below the range of the first quartile plus three interquartile ranges or above the third quartile plus three interquartile ranges of the data. The LMER models for visual tetanization are reported in Table 4. Changes in P1 peak amplitude following visual tetanization interacted with age group. Post-hoc comparisons found that younger adults exhibited a marginal decrease in P1 peak amplitude following visual tetanization (p<0.1), but older adults did not. N1 peak amplitudes decreased (became more positive) following visual tetanization across age groups. N1b exhibited a marginal decrease in amplitude following visual tetanization across age groups (p<0.1). C1, P2a, and P2 exhibited no change in amplitude following visual tetanization. Adding tetanization condition (horizontal or vertical tetanization) and stimulus type (horizontal or vertical gradient), along with their interaction terms with trial and age group, did not significantly improve model fit (p>0.1), suggesting that changes in response amplitudes following tetanization were not specific to the tetanizing stimulus. Adding visual acuity (Snellen) and contrast sensitivity (Pelli-Robson) scores to these models did not change the pattern of significant relationships or improve model fit (p>0.1).

**Table 4.**
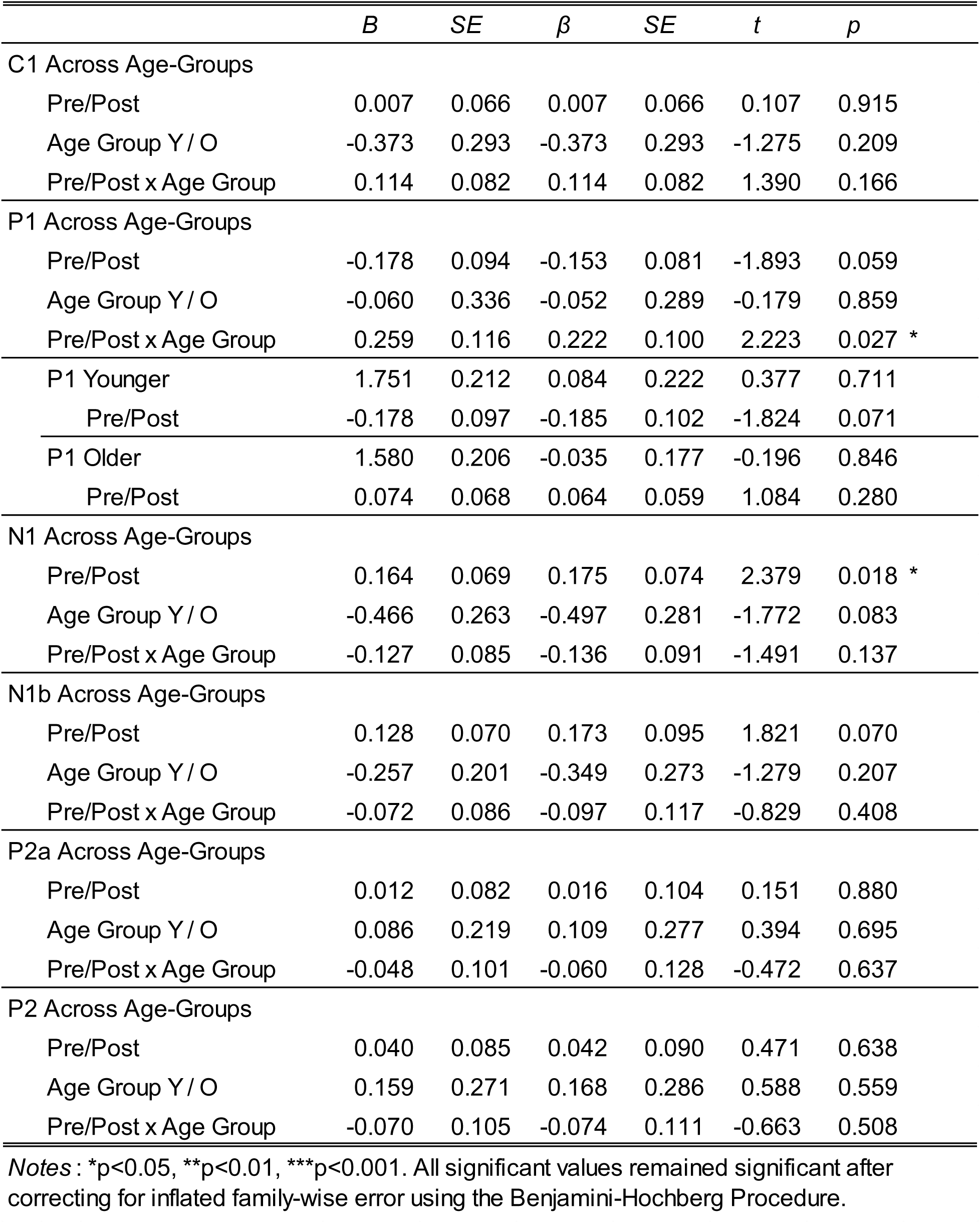
Visual Tetanization Linear Mixed Effects Models of N1 Peak Ampitude Predicted by Trial and Age Group

#### Summary

The results of our tetanization study find that the N1 and P2 peak amplitudes of auditory-evoked ERPs can decrease following auditory tetanization. Decreases in the N1 amplitudes of auditory-evoked potentials were smaller for older adults than for younger adults, the first such report of an age-group difference in modulation of auditory-evoked potentials after tetanization. Similarly, we found that the N1 peak amplitudes of visual-evoked ERPs can also decrease following visual tetanization, but these decreases in N1 peak amplitudes were similar across age groups. Though we did find a significant interaction between the effects of visual tetanization and age-group on the visual-evoked P1, multiple comparisons found that that neither age group exhibited a significant change in P1 amplitude following visual tetanization.

The decreases in response amplitudes that we observed following auditory and visual tetanization were not specific to the tetanized stimulus. Tetanization to either 1 kHz tone bursts or 4 kHz tone bursts resulted in decreases in the response amplitudes of potentials evoked by either tone stimulus. Similarly, tetanization to either the horizontal or vertical sine gradient resulted in decreases in the response amplitudes of potentials evoked by either sine gradient. Importantly, the decreases in N1 and P2 amplitude after auditory tetanization were not significantly different from the decreases in N1 and P2 amplitude following slow presentation of 1 kHz and 4 kHz tone bursts, suggesting that high-frequency stimulus presentation is not needed to modulate the amplitude of auditory evoked potentials.

Our results join a growing number of studies reporting contrasting effects of sensory tetanization on sensory-evoked potentials. To determine the extent to which sensory tetanization effects are significant across studies, we conducted meta-analyses.

## Meta-Analyses

Our finding a decrease in P1 and N1 peak amplitude after tetanization is consistent with the findings of some studies, but not others. Our study joins numerous others that have used sensory tetanization to elicit changes in ERPs. While sensory tetanization may be a valuable non-invasive approach to induce LTP-like plasticity in humans (Kirk et al., 2021; Sumner, Spriggs, et al., 2020), contrasting effects across studies must be considered. As previously discussed, some studies find that tetanization can increase the amplitude of event related potentials (Kleeva et al., 2022; Lei et al., 2017; Lengali et al., 2021; McNair et al., 2006; Moore et al., 2020; Ross et al., 2008; Rygvold et al., 2020; Smallwood et al., 2015; Spriggs et al., 2017; Spriggs et al., 2018; Spriggs et al., 2019; Sumner et al., 2018; Wilson et al., 2017), others find no change in response amplitude following tetanization (D’Souza et al., 2018; Normann et al., 2007; Rygvold et al., 2020; Sumner et al., 2018), and still others, including ourselves, find that response amplitudes decrease after tetanization (Forsyth et al., 2015; Kleeva et al., 2022; Klöppel et al., 2015; Rebreikina et al., 2021; Spriggs et al., 2018; Wynn et al., 2019). These conflicting results are troubling, especially when considering that many studies use methods similar to those initially reported by Clapp, Kirk, et al. (2005) and Teyler et al. (2005). Recent reviews of the growing number of studies employing tetanization as a method for studying cortical plasticity have not systematically examined the contrasting effects of tetanization reported across studies (Kirk et al., 2021; Sanders et al., 2018; Sumner, Spriggs, et al., 2020). Here, we conducted meta-analyses to determine whether the average effect of sensory tetanization is significant and, subsequently, whether tetanization is a reliable approach to induce LTP-like plasticity in the human brain.

### Methods

#### Studies

We first conducted a citation search in Google Scholar to find papers that cite either Clapp, Kirk, et al. (2005) or Teyler et al. (2005) These two seminal papers introduced the sensory tetanization paradigm and we reasoned that subsequent papers using the paradigm would site at least one of them. Search results were then screened for empirical studies that used “high frequency stimulation” or “sensory tetanization” to modulate responses to sensory stimuli, resulting in a list of 42 studies (including our own tetanization study reported here). From this list, we excluded 6 studies that did not employ EEG to evaluate changes in sensory-evoked ERPs after tetanization. We then excluded 8 studies that did not parallel the original approach of Clapp, Kirk, et al. (2005) or Teyler et al. (2005) and conduct ERP analyses that explicitly test for amplitude changes in the components of sensory evoked potentials. We further excluded 6 studies that lacked a group of younger healthy participants. One other study (Moore & Loprinzi, 2021) was excluded for reporting data overlapping with a previous study (Moore et al., 2020) to avoid any explicit repetition of study findings. The remaining 21 studies were divided based on their study of visual tetanization (n=16) or auditory tetanization (n=6), the details of which are reported in Table 5 and Table 6, respectively. Studies that did not fit the criteria for our meta-analyses still report findings that are important for understanding the reliability of sensory tetanization effects and are considered in the Discussion.

**Table 5.**
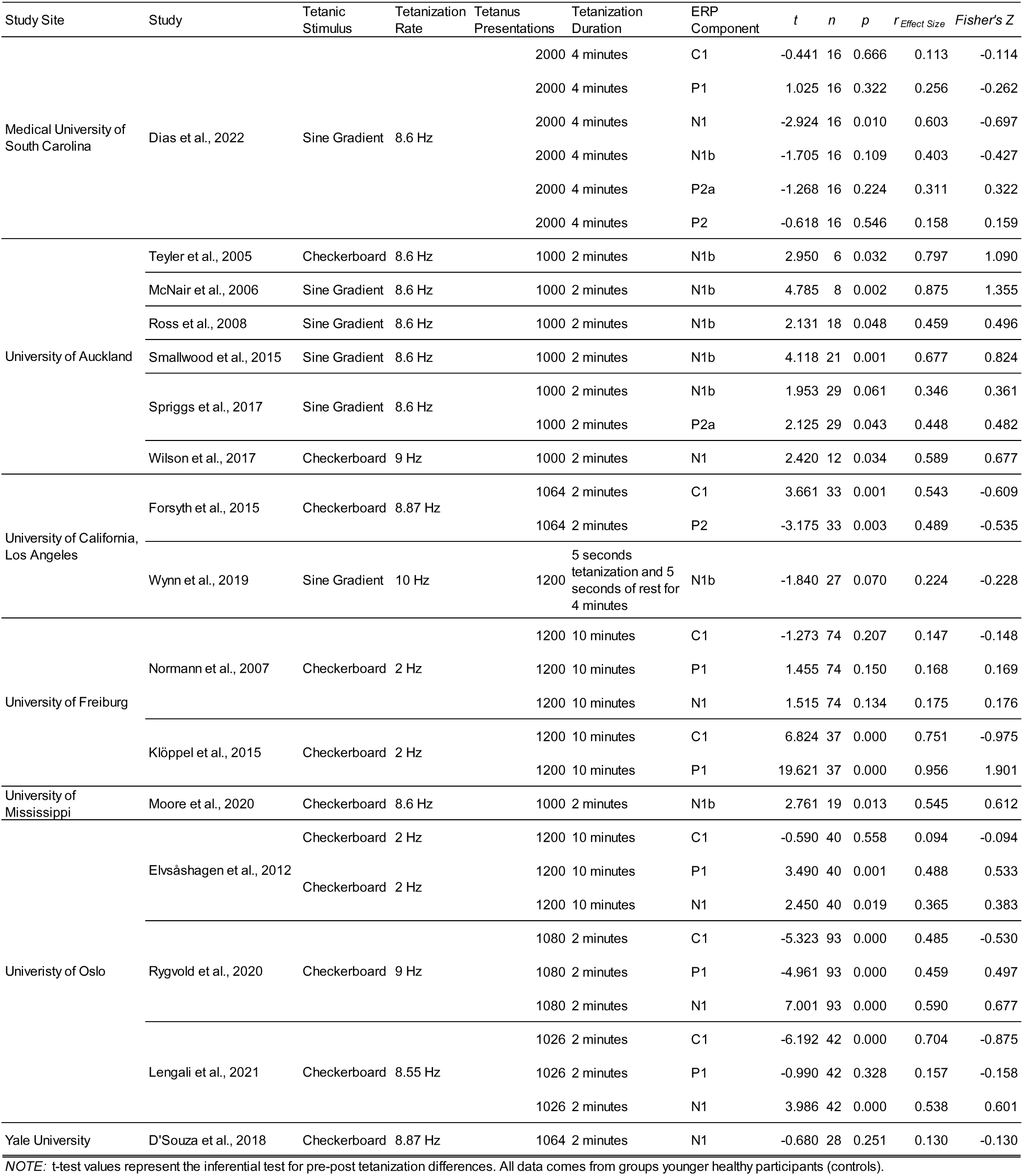
Summary of studies of visual tetanization included in the meta-analysis

**Table 6.**
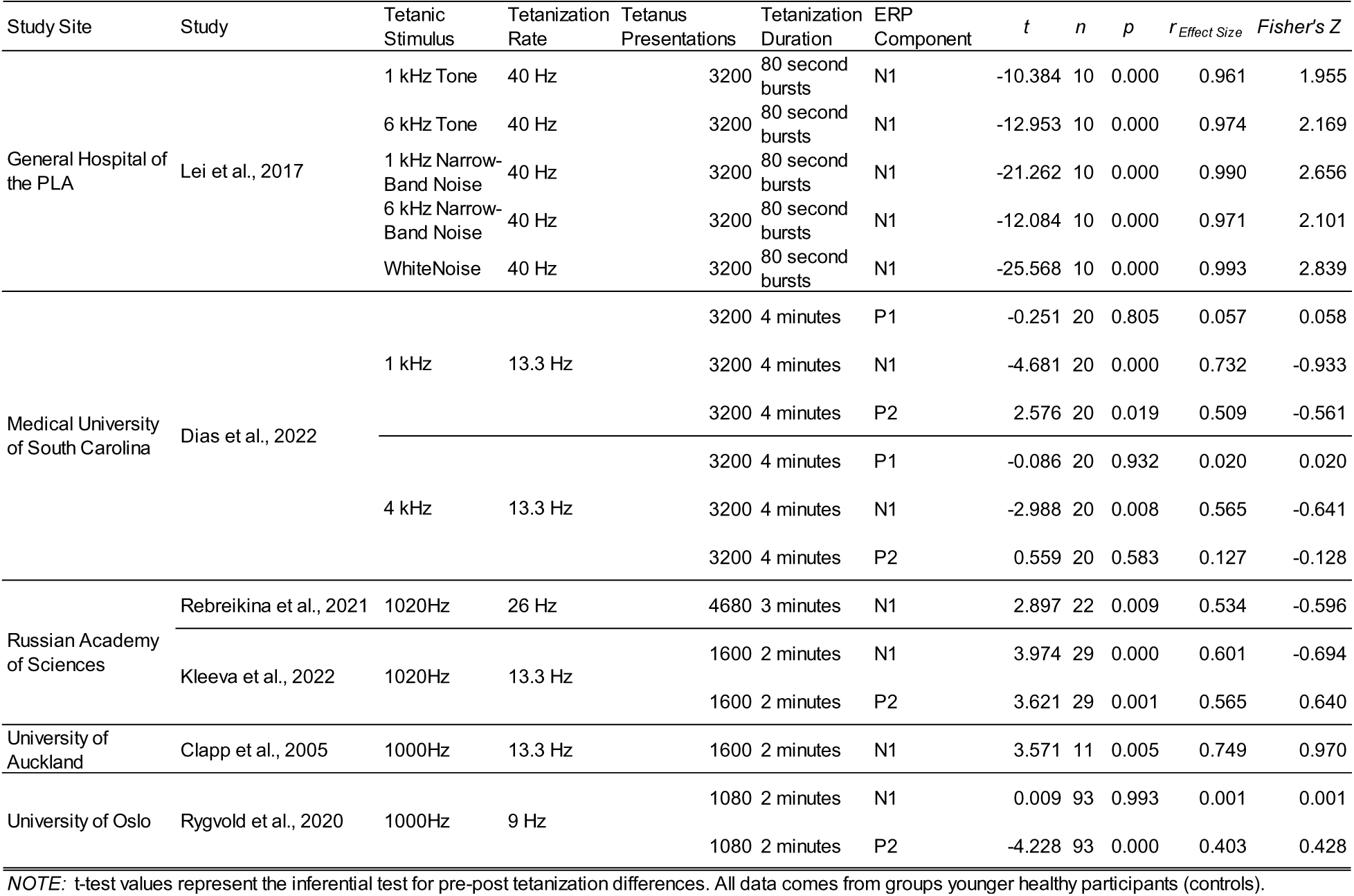
Summary of studies of auditory tetanization included in the meta-analysis

#### Analytical Approach

For each of the 21 studies included in our meta-analysis, we identified the reported ERP component(s) and computed the effect size (r_Effect Size_) for each component’s change in amplitude following tetanization. Effect sizes were computed from the descriptive and inferential statistics available for the group of younger healthy adults reported in each study. The effect sizes for each component reported in each study were then transformed into Z scores using Fisher’s transformation and were then coded as reflecting an increase (positive value) or decrease (negative value) in amplitude following tetanization. These Z scores were used as the outcome variable of our meta-analyses (e.g., Hansen et al., 2022; Rosenthal, 1991; Rosnow et al., 2000).

Many studies report multiple ERP components for multiple tetanization conditions, where participants were tetanized and tested on different stimuli, like in our own study (e.g., tetanization to 1 kHz and 4 kHz tones). In such cases, different components and tetanization conditions may be nested within a single study. To account for the variability between components and conditions nested within studies, LMER was used to conduct our meta-analyses. Different LMER models were used to perform separate meta-analyses for studies of visual tetanization and for studies of auditory tetanization. These LMER models included the Z transforms of effect sizes as the outcome variable and study as a random factor (grouping variable). Each study was weighted by their reported sample size. Component was included in these models as a fixed factor along with a zero-intercept parameter to determine the degree to which the average effect sizes of each component differed from zero.

The studies included in these study-level meta-analyses were each conducted at one of nine study sites. As such, many of these studies were conducted at the same study site (by the same research group), yet these studies do not provide enough detail regarding their sample composition to know whether participants’ tetanization data overlapped across different studies. To control for this uncertainty, we conducted follow-up meta-analyses that nested studies within study sites, including both study and study site as random factors (grouping variables).

Our meta-analyses were performed using the *meta* (Balduzzi et al., 2019) and *metafor* (Viechtbauer & Cheung, 2010) packages for *R* (Harrer et al., 2021; R Core Team, 2022).

### Results

Table 7 and Figure 9 report the results of our meta-analysis of studies of visual tetanization. Across studies, the only visual-evoked ERP component that appears to demonstrate a consistent change in response amplitude following visual tetanization is C1. Across studies, C1 exhibits a significant amplitude decrease (becomes less negative) following visual tetanization. Importantly, when studies are nested within study sites, no component of the visual-evoked ERP demonstrates a consistent change in response amplitude following visual tetanization.

**Table 7.**
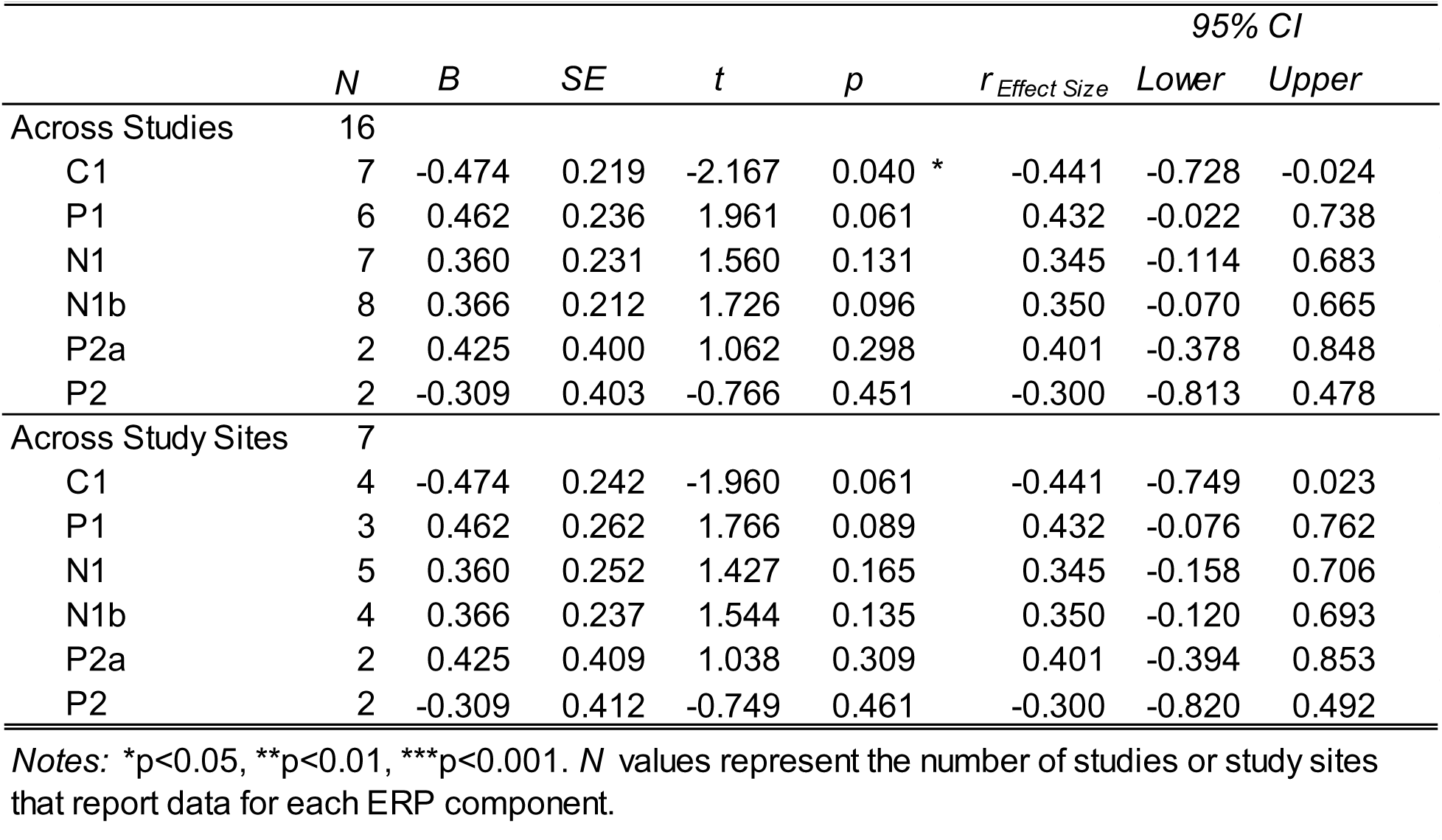
Meta analysis of visual tetanization effects for each ERP component tested across studies and across study sites

**Figure 9.**
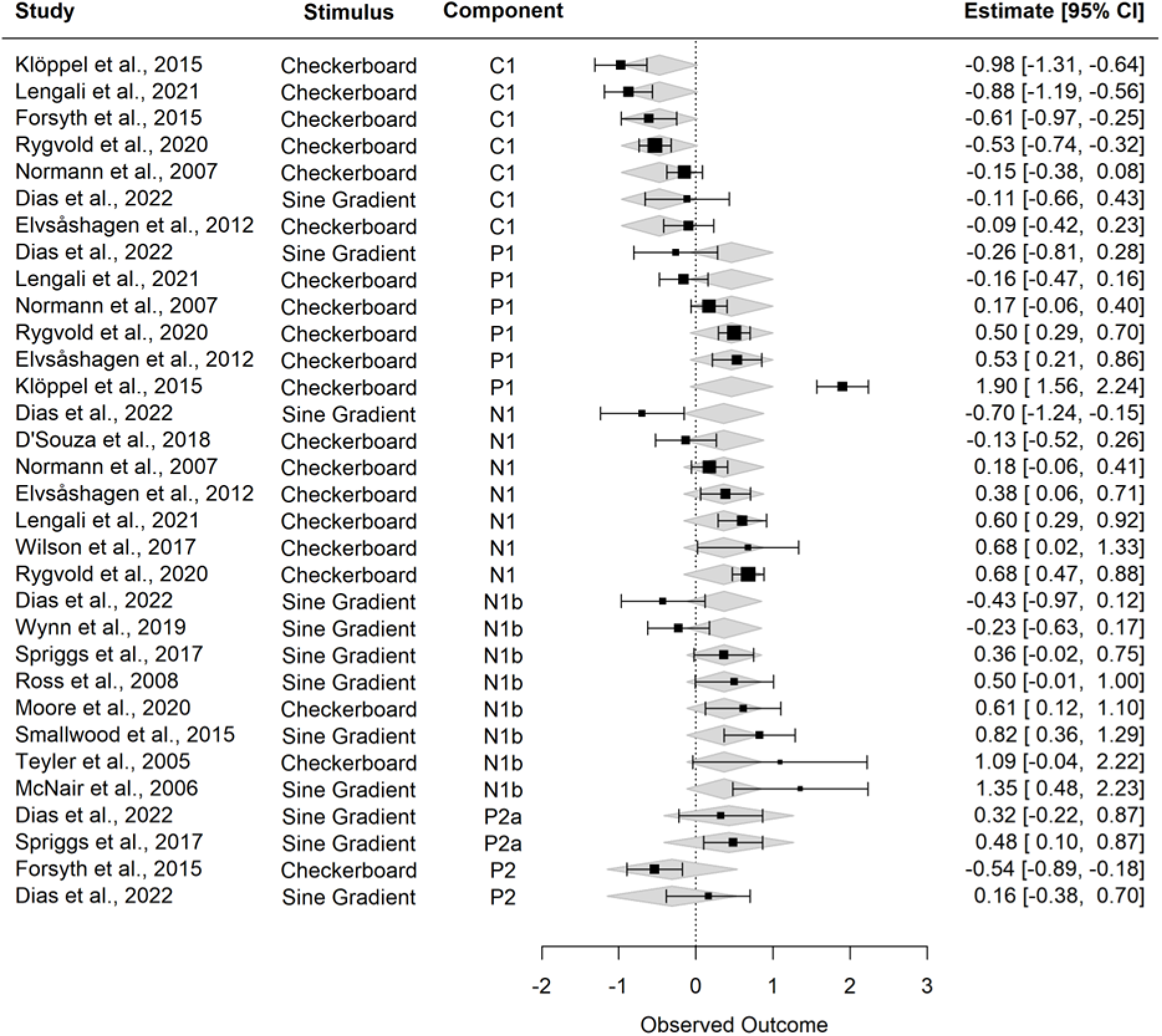
Forest plot of our meta-analysis of visual tetanization studies. Study data are organized by visual-evoked ERP component, though ERP components may be nested within individual studies. Boxes and whiskers represent the Z-transformed effect size and 95% confidence interval for each study (reported numerically on the right). Box size represents the weight of each study, determined by study sample size. Individual study data for each ERP component are reported with the average Z-transformed effect size and 95% confidence interval for each component across studies (grey diamonds).

Table 8 and Figure 10 report the results of our meta-analysis of studies of auditory tetanization. Across studies, no auditory-evoked ERP component demonstrates a consistent change in response amplitude following auditory tetanization. Nesting studies within study sites did not change this pattern of results.

**Table 8.** Meta analysis of auditory tetanization effects for each ERP component tested across studies and across study sites

|  | *N* | *B* | *SE* | *t* | *p* | *r* *Effect Size* | 95% *CI* |  |
| --- | --- | --- | --- | --- | --- | --- | --- | --- |
|  |  |  |  |  |  |  | *Lower* | *Upper* |
| Across Studies | 6 |  |  |  |  |  |  |  |
| P1 | 1 | 0.039 | 1.222 | 0.032 | 0.975 | 0.039 | -0.989 | 0.990 |
| N1 | 6 | 0.258 | 0.595 | 0.434 | 0.671 | 0.252 | -0.769 | 0.911 |
| P2 | 3 | 0.275 | 0.795 | 0.346 | 0.735 | 0.268 | -0.892 | 0.963 |
| Across Study Sites | 5 |  |  |  |  |  |  |  |
| P1 | 1 | 0.039 | 1.308 | 0.030 | 0.977 | 0.039 | -0.992 | 0.993 |
| N1 | 5 | 0.258 | 0.664 | 0.389 | 0.703 | 0.252 | -0.823 | 0.933 |
| P2 | 3 | 0.275 | 0.851 | 0.323 | 0.751 | 0.268 | -0.914 | 0.970 |

*Notes:* \* $p < 0.05$ , \*\* $p < 0.01$ , \*\*\* $p < 0.001$ . *N* values represent the number of studies or study sites that report data for each ERP component.

**Figure 10.**
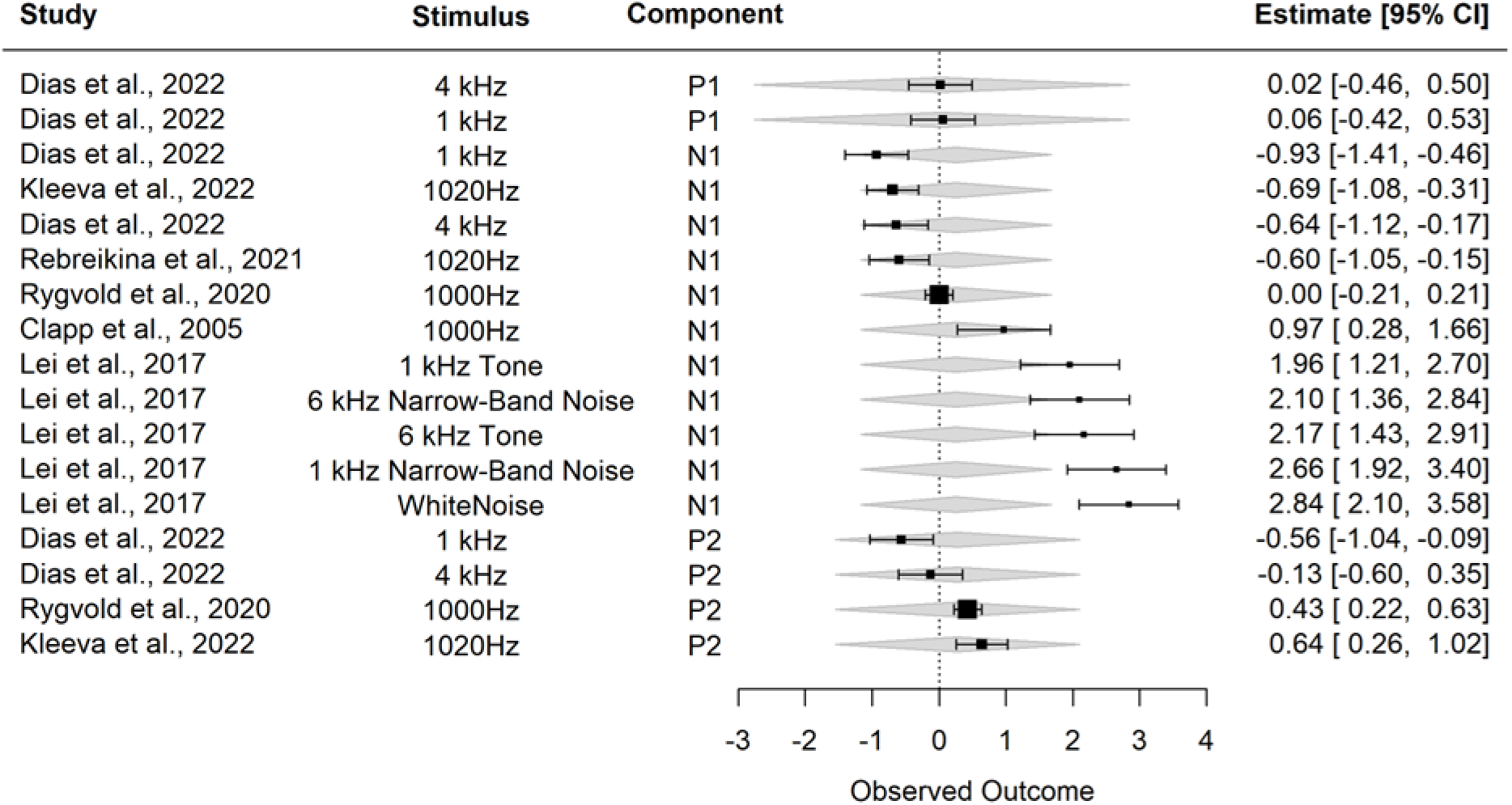
Forest plot of our meta-analysis of auditory tetanization studies. Study data are organized by auditory-evoked ERP component, though ERP components may be nested within individual studies. Boxes and whiskers represent the Z-transformed effect size and 95% confidence interval for each study (reported numerically on the right). Box size represents the weight of each study, determined by study sample size. Individual study data for each ERP component are reported with the average Z-transformed effect size and 95% confidence interval for each component across studies (grey diamonds).

### Summary

Our meta-analyses suggest that sensory tetanization does no produce reliable changes in cortical response amplitudes. Across both visual and auditory studies, no single component produced a significant amplitude increase reflective of LTP-like plasticity. Some components of the visual-evoked and auditory-evoked ERP have been studied more than others. More studies report effects of visual tetanization on the visual C1, P1, N1, and N1b than on the visual P2a and P2. Similarly, more studies report effects of auditory tetanization on the auditory N1 than on the auditory P1 or P2. Regardless of this fact, the average effects of sensory tetanization on those components that have been studied more are not significant across studies.

Evident from Table 5 and Table 6 is substantial variability in the methods for conducting sensory tetanization studies. Differences in stimuli, presentation rate, and tetanization duration may all contribute to the variable effects of tetanization between studies, visible in Figure 9 and Figure 10. Variability between study methods and study results begs the question of how many studies employing traditional and experimental approaches to sensory tetanization produce null results and are never reported. Our meta-analyses included only published studies, subject to publication bias, but consideration should be given to “file-drawer” studies that never make it to publication (Rosenthal, 1979; Rosenthal, 1991). Future studies should consider how different experimental approaches to tetanization may produce different physiological effects.

## Discussion

There is much interest in developing non-invasive approaches to study cortical plasticity in the human brain, but contrasting results across studies suggest that sensory tetanization (Clapp, Kirk, et al., 2005; Teyler et al., 2005) may not be a reliable experimental approach in humans. Our study of auditory tetanization found that N1 and P2 peak amplitudes decreased after tetanization. These amplitude decreases were not specific to the tetanizing stimulus, such that tetanization to either the 1 kHz or 4 kHz tone burst decreased the N1 and P2 peak amplitudes of responses to both stimuli. We found that older adults exhibited a more modest decrease in N1 peak amplitude after tetanization, but these effects of tetanization were no different from our slow-presentation control across age groups. The results of our study of visual tetanization were similar to our auditory findings, with participants exhibiting decreases in N1 amplitude after visual tetanization that were not stimulus specific. However, unlike our study of auditory tetanization, these decreases in visual N1 amplitude were similar for younger and older adults. Our auditory and visual tetanization results contribute to the growing number of contrasting effects reported across studies of sensory tetanization. Our meta-analyses of a subset of these studies found that the average effect of visual and auditory tetanization on sensory-evoked ERPs are modest at best or otherwise non-significant. Variability in study methodology may account for some of the contrasting effects of sensory tetanization on ERP amplitudes, but other inconsistent and conflicting findings concerning age effects and stimulus specificity raise important concerns regarding the validity of sensory tetanization as a tool for studying human cortical plasticity in vivo.

### Methodological Variability and Tetanization Effects

As previously discussed, methodological differences between studies may account for some variability in tetanization effects found in our meta-analyses, but contrasting effects of sensory tetanization are not limited to those studies. As stated earlier, fMRI studies report BOLD increases (Clapp, Zaehle, et al., 2005; Wijtenburg et al., 2017; Zaehle et al., 2007) and decreases (Lahr et al., 2014) after sensory tetanization. Similarly, visual studies concerned primarily with localizing the source of N1 and N1b amplitude changes after sensory tetanization report amplitude increases (Spriggs et al., 2019), decreases (Spriggs et al., 2018), and even no change in amplitude (Sumner et al., 2018). In fact, two EEG studies not included in our meta-analyses for reporting global metrics of response amplitude (no individual components were measured) found that overall ERP amplitudes decreased after visual tetanization (Abuleil et al., 2019) and after auditory tetanization (Mears & Spencer, 2012).

The contrasting effects of sensory tetanization that are observed across visual and auditory studies suggest that sensory tetanization may not be a reliable experimental tool for studying LTP-like plasticity. Similar tetanization paradigms should not produce amplitude increases attributed to LTP in some studies (Kirk et al., 2021; Sanders et al., 2018; Sumner, Spriggs, et al., 2020) and amplitude decreases sometimes attributed to long-term depression (LTD, the opposite of LTP) in others (e.g., Abuleil et al., 2019; Kleeva et al., 2022). Methodological differences between studies may be able to account for some of this variability in tetanization effects. If this is the case, however, it would suggest that tetanization effects are highly sensitive to experimental paradigm and analytical approach, raising questions regarding the generalizability of the results from one study to another and the degree to which tetanization effects reflect LTP-like plasticity in the human brain. Meaningful inference of the mechanisms that underlie changes in sensory-evoked response amplitudes after sensory tetanization is difficult given the contrasting effects reported across studies, especially if these contrasting effects reflect (at least in part) the sensitivity of tetanization effects to specific study methodologies. Future studies should investigate how tetanization affects are modulated by stimulus type, presentation rate, and tetanization duration to better understand the physiological changes that follow tetanization and how they reflect neuroplastic changes in cortical function.

It is also important to consider our finding that changes in N1 amplitude after auditory tetanization do not differ from changes in N1 amplitude after slow stimulus presentation, further complicating interpretation of current and past results and the role of experimental paradigm on tetanization effects. Additionally, the contrasting effects of tetanization on ERP amplitudes are only one example of the conflicting results found across tetanization studies.

### Age Effects on Sensory Tetanization

We found that older adults exhibited a significant decrease in the N1 amplitudes of auditory-evoked and visual-evoked potentials after auditory and visual tetanization, respectively. Finding that older adults can exhibit a change in sensory-evoked response amplitudes after sensory tetanization is inconsistent with studies finding that older adults do not demonstrate *visual*-evoked response amplitude changes after *visual* tetanization (Abuleil et al., 2019; Spriggs et al., 2017). There is one study that reported a change in visual-evoked response amplitudes in older adults, but the response amplitudes *increased* after visual tetanization, contrasting with our results (de Gobbi Porto et al., 2015). Our finding that older adults demonstrated a smaller decrease in *auditory*-evoked response amplitudes than younger adults after auditory tetanization is consistent with one study finding similar age-group differences in the decrease in *visual*-evoked amplitudes after visual tetanization (Abuleil et al., 2019), but it is inconsistent with another study finding that age is unrelated to changes in *visual*-evoked response amplitudes after visual tetanization (Valstad et al., 2020). In contrast, our finding that decreases in *visual*-evoked response amplitudes after visual tetanization are similar for younger and older adults is inconsistent with the one study finding older adults exhibit a smaller decrease in *visual*-evoked response amplitudes after visual tetanization (Abuleil et al., 2019), but it is consistent with the study finding no relationship between age and visual tetanization (Valstad et al., 2020). Finding that the effects of auditory tetanization are weaker for older adults has not been reported previously. It is tempting to infer a relationship between age and the mechanisms that underly auditory plasticity from these results, as others have done when studying age-group differences in visual tetanization (Abuleil et al., 2019; Spriggs et al., 2017), but the inconsistencies across our results and others’ described above raise concerns regarding the reliability and validity of such age effects when using sensory tetanization to study age-related differences in neural plasticity.

### Stimulus Specificity of Sensory Tetanization

We found no evidence of stimulus-specific changes in sensory-evoked response amplitudes after sensory tetanization. Tetanization to either the 1 kHz or 4 kHz tone similarly decreased the auditory-evoked response amplitudes of both the 1 kHz and 4 kHz tones. Similarly, tetanization to either the horizontal or vertical sine gradient similarly decreased the visual-evoked response amplitudes of both the horizontal and vertical sign gradients. Other studies have failed to find consistent stimulus-specific changes in sensory evoked responses after similar auditory tone-burst tetanization (Kleeva et al., 2022; Rebreikina et al., 2021) and visual sine-gradient tetanization (Cooke & Bear, 2010; Sumner et al., 2018). However, several other studies have found changes in sensory-evoked responses that are specific to the tetanized auditory tone-burst (Kompus & Westerhausen, 2018; Mears & Spencer, 2012) and visual sine-gradient (McNair et al., 2006; Ross et al., 2008; Spriggs et al., 2017; Wynn et al., 2019). The results across studies suggest that tetanization effects may sometimes affect mechanisms that generalize across stimulus types. What determines when sensory tetanization effects are sometimes stimulus-specific is unclear and should be considered in future studies.

### Conclusion

Our study of auditory and visual tetanization joins a large body of work with highly variable findings and contrasting results. Changes in auditory-evoked and visual-evoked response amplitudes after sensory tetanization contrast across different studies. Studies testing the extent to which tetanization effects are modulated by age or whether tetanization effects are specific to the tetanizing stimulus have also yielded inconsistent results. Variability is also found in how long tetanizing effects last and whether tetanization effects relate to behavior (for reviews, see Kirk et al., 2021; Sanders et al., 2018; Sumner, Spriggs, et al., 2020). Though we did not discuss these last two topics in detail, they contribute to the overall lack of reliable results evident across sensory tetanization studies.

Variable effects between studies are inevitable. Aside from the variability attributed to different researchers, equipment, and samples of participants (just to name a few), methods are often manipulated to test new hypotheses and advance understanding of a mechanism. Methodological differences in tetanization paradigms and analytical approaches may be able to partially account for the variable results observed across tetanization studies. Future work should investigate this possibility and examine how they may affect the generalizability of results from one study to another and to what extent tetanization-induced changes in visual-evoked and auditory-evoked responses reflect LTP-like plasticity.

## Sources of Funding

This work was supported (in part) by grants from the National Institute on Deafness and Other Communication Disorders (NIDCD) (R01 DC014467, R01 DC017619, P50 DC000422, and T32 DC014435) and the Hearing Health Foundation. The project also received support from the South Carolina Clinical and Translational Research (SCTR) Institute with an academic home at the Medical University of South Carolina, National Institute of Health/National Center for Research Resources (NIH/NCRR) Grant number UL1 TR001450. This investigation was conducted in a facility constructed with support from Research Facilities Improvement Program Grant Number C06 RR14516 from the National Center for Research Resources, NIH. The authors declare no competing financial interests.

## Abbreviations

BOLD: blood-oxygen level dependent
EEG: electroencephalography
ERP: event-related potential
fMRI: functional magnetic resonance imaging
LMER: linear mixed-effects model
LTP: long-term potentiation
rTMS: repeated transcranial magnetic stimulation

## Conflict of Interest Statement

The authors declare no competing financial interests.

## Data Sharing Statement

Deidentified data are available upon request

## Credit Author Statement

**James W. Dias:** Conceptualization, Methodology, Formal Analysis, Investigation, Data Curation, Writing – Original Draft, Visualization. **Carolyn M. McClaskey:** Investigation, Data Curation, Writing – Review & Editing. **Jeffrey A. Rumschlag:** Investigation, Data Curation, Writing – Review & Editing. **Kelly C. Harris:** Conceptualization, Methodology, Investigation, Resources, Data Curation, Writing – Review & Editing, Supervision, Project Administration, Funding Acquisition.

## Notes

### Competing Interest Statement

The authors have declared no competing interest.

### Summary of Updates

The revised manuscript incorporates changes to text to improve clarity and additional analyses to further support the original outcomes.

## References

Abuleil, D., McCulloch, D. L., & Thompson, B. (2019). Older Adults Exhibit Greater Visual Cortex Inhibition and Reduced Visual Cortex Plasticity Compared to Younger Adults [Original Research]. Frontiers in Neuroscience, 13. https://doi.org/10.3389/fnins.2019.00607

Anderer, P., Semlitsch, H. V., & Saletu, B. (1996). Multichannel auditory event-related brain potentials: effects of normal aging on the scalp distribution of N1, P2, N2 and P300 latencies and amplitudes. Electroencephalography and Clinical Neurophysiology, 99(5), 458–472. https://doi.org/10.1016/S0013-4694(96)96518-9

Balduzzi, S., Rücker, G., & Schwarzer, G. (2019). How to perform a meta-analysis with R: a practical tutorial. Evidence Based Mental Health, 22(4), 153–160. https://doi.org/10.1136/ebmental-2019-300117

Beck, H., Goussakov, I. V., Lie, A., Helmstaedter, C., & Elger, C. E. (2000). Synaptic Plasticity in the Human Dentate Gyrus. The Journal of Neuroscience, 20(18), 7080–7086. https://doi.org/10.1523/jneurosci.20-18-07080.2000

Bliss, T. V., & Lømo, T. (1973). Long-lasting potentiation of synaptic transmission in the dentate area of the anaesthetized rabbit following stimulation of the perforant path. The Journal of Physiology, 232(2), 331–356. https://doi.org/10.1113/jphysiol.1973.sp010273

Bliss, T. V. P., & Collingridge, G. L. (1993). A synaptic model of memory: long-term potentiation in the hippocampus. Nature, 361(6407), 31–39. https://doi.org/10.1038/361031a0

Çavuş, I., Reinhart, R. M. G., Roach, B. J., Gueorguieva, R., Teyler, T. J., Clapp, W. C., Ford, J. M., Krystal, J. H., & Mathalon, D. H. (2012). Impaired Visual Cortical Plasticity in Schizophrenia. Biological Psychiatry, 71(6), 512–520. https://doi.org/10.1016/j.biopsych.2012.01.013

Chen, W. R., Lee, S., Kato, K., Spencer, D. D., Shepherd, G. M., & Williamson, A. (1996). Long-term modifications of synaptic efficacy in the human inferior and middle temporal cortex. Proceedings of the National Academy of Sciences, 93(15), 8011–8015. https://doi.org/10.1073/pnas.93.15.8011

Chung, S. W., Hill, A. T., Rogasch, N. C., Hoy, K. E., & Fitzgerald, P. B. (2016). Use of theta-burst stimulation in changing excitability of motor cortex: A systematic review and meta-analysis. Neuroscience & Biobehavioral Reviews, 63, 43–64. https://doi.org/10.1016/j.neubiorev.2016.01.008

Clapp, W. C., Kirk, I. J., Hamm, J. P., Shepherd, D., & Teyler, T. J. (2005). Induction of LTP in the human auditory cortex by sensory stimulation. European Journal of Neuroscience, 22(5), 1135–1140. doi: https://doi.org/10.1111/j.1460-9568.2005.04293.x

Clapp, W. C., Zaehle, T., Lutz, K., Marcar, V. L., Kirk, I. J., Hamm, J. P., Teyler, T. J., Corballis, M. C., & Jancke, L. (2005). Effects of long-term potentiation in the human visual cortex: a functional magnetic resonance imaging study. NeuroReport, 16(18). https://journals.lww.com/neuroreport/Fulltext/2005/12190/Effects_of_long_term_potentiation_in_the_human.1.aspx

Clogg, C. C., Petkova, E., & Haritou, A. (1995). Statistical Methods for Comparing Regression Coefficients Between Models. American Journal of Sociology, 100(5), 1261–1293. https://doi.org/10.1086/230638

Cohen, J., Cohen, P., West, S. G., & Aiken, L. S. (2003). Applied Multiple Regression/Correlation Analysis for the Behavioral Sciences. Lawrency Erlbaum Associates. https://doi.org/10.4324/9781410606266

Cooke, S. F., & Bear, M. F. (2010). Visual Experience Induces Long-Term Potentiation in the Primary Visual Cortex. The Journal of Neuroscience, 30(48), 16304–16313. https://doi.org/10.1523/jneurosci.4333-10.2010

Cooke, S. F., & Bliss, T. V. P. (2006). Plasticity in the human central nervous system. Brain, 129(7), 1659–1673. https://doi.org/10.1093/brain/awl082

D’Souza, D. C., Carson, R. E., Driesen, N., Johannesen, J., Ranganathan, M., Krystal, J. H., Ahn, K.-H., Bielen, K., Carbuto, M., Deaso, E., D’Souza, D. C., Ranganathan, M., Naganawa, M., Ranganathan, M., D’Souza, D. C., Nabulsi, N., Zheng, M.-Q., Lin, S.-f., Huang, Y., … Pittman, B. (2018). Dose-Related Target Occupancy and Effects on Circuitry, Behavior, and Neuroplasticity of the Glycine Transporter-1 Inhibitor PF-03463275 in Healthy and Schizophrenia Subjects. Biological Psychiatry, 84(6), 413–421. https://doi.org/10.1016/j.biopsych.2017.12.019

de Gobbi Porto, F. H., Fox, A. M., Tusch, E. S., Sorond, F., Mohammed, A. H., & Daffner, K. R. (2015). In vivo evidence for neuroplasticity in older adults. Brain Research Bulletin, 114, 56–61. https://doi.org/10.1016/j.brainresbull.2015.03.004

Delorme, A., & Makeig, S. (2004). EEGLAB: an open source toolbox for analysis of single-trial EEG dynamics including independent component analysis. Journal of Neuroscience Methods, 134(1), 9–21. https://doi.org/10.1016/j.jneumeth.2003.10.009

Dias, J. W., McClaskey, C. M., & Harris, K. C. (2018). Time-compressed speech identification is predicted by auditory neural processing, perceptuomotor speed, and executive functioning in younger and older listeners. Journal of the Association for Research in Otolaryngology. https://doi.org/10.1007/s10162-018-00703-1

Dias, J. W., McClaskey, C. M., & Harris, K. C. (2021). Early auditory cortical processing predicts auditory speech in noise identification and lipreading. Neuropsychologia, 161, 108012. https://doi.org/10.1016/j.neuropsychologia.2021.108012

Elvsåshagen, T., Moberget, T., Bøen, E., Boye, B., Englin, N. O. A., Pedersen, P. Ø., Andreassen, O. A., Dietrichs, E., Malt, U. F., & Andersson, S. (2012). Evidence for Impaired Neocortical Synaptic Plasticity in Bipolar II Disorder. Biological Psychiatry, 71(1), 68–74. https://doi.org/10.1016/j.biopsych.2011.09.026

Esser, S. K., Huber, R., Massimini, M., Peterson, M. J., Ferrarelli, F., & Tononi, G. (2006). A direct demonstration of cortical LTP in humans: A combined TMS/EEG study. Brain Research Bulletin, 69(1), 86–94. https://doi.org/10.1016/j.brainresbull.2005.11.003

Folstein, M. F., Robins, L. N., & Helzer, J. E. (1983). The Mini-Mental State Examination. Archives of General Psychiatry, 40(7), 812. https://doi.org/10.1001/archpsyc.1983.01790060110016

Forsyth, J. K., Bachman, P., Mathalon, D. H., Roach, B. J., & Asarnow, R. F. (2015). Augmenting NMDA receptor signaling boosts experience-dependent neuroplasticity in the adult human brain. Proceedings of the National Academy of Sciences, 112(50), 15331–15336. https://doi.org/10.1073/pnas.1509262112

Forsyth, J. K., Bachman, P., Mathalon, D. H., Roach, B. J., Ye, E., & Asarnow, R. F. (2017). Effects of Augmenting N-Methyl-D-Aspartate Receptor Signaling on Working Memory and Experience-Dependent Plasticity in Schizophrenia: An Exploratory Study Using Acute d-cycloserine. Schizophrenia Bulletin, 43(5), 1123–1133. https://doi.org/10.1093/schbul/sbw193

Hamilton, H. K., Roach, B. J., Cavus, I., Teyler, T. J., Clapp, W. C., Ford, J. M., Tarakci, E., Krystal, J. H., & Mathalon, D. H. (2020). Impaired Potentiation of Theta Oscillations During a Visual Cortical Plasticity Paradigm in Individuals With Schizophrenia [Original Research]. Frontiers in Psychiatry, 11. https://doi.org/10.3389/fpsyt.2020.590567

Hansen, C., Steinmetz, H., & Block, J. (2022). How to conduct a meta-analysis in eight steps: a practical guide. Management Review Quarterly, 72(1), 1–19. https://doi.org/10.1007/s11301-021-00247-4

Harrer, M., Cuijpers, P., Furukawa, T. A., & Ebert, D. D. (2021). Doing Meta-Analysis with R: A Hands-On Guide. Chapman and Hall/CRC. https://doi.org/10.1201/9781003107347

Harris, K. C., Wilson, S., Eckert, M. A., & Dubno, J. R. (2012). Human Evoked Cortical Activity to Silent Gaps in Noise: Effects of Age, Attention, and Cortical Processing Speed. Ear and Hearing, 33(3), 330–339. https://doi.org/10.1097/AUD.0b013e31823fb585

Jahshan, C., Wynn, J. K., Mathalon, D. H., & Green, M. F. (2017). Cognitive correlates of visual neural plasticity in schizophrenia. Schizophrenia Research, 190, 39–45. https://doi.org/10.1016/j.schres.2017.03.016

Kandel, E. R., & Tauc, L. (1964). Mechanism of Prolonged Heterosynaptic Facilitation. Nature, 202(4928), 145–147. https://doi.org/10.1038/202145a0

Kandel, E. R., & Tauc, L. (1965). Heterosynaptic facilitation in neurones of the abdominal ganglion of Aplysia depilans [https://doi.org/10.1113/jphysiol.1965.sp007742]. The Journal of Physiology, 181(1), 1–27. https://doi.org/10.1113/jphysiol.1965.sp007742

Kirk, I. J., Spriggs, M. J., & Sumner, R. L. (2021). Human EEG and the mechanisms of memory: investigating long-term potentiation (LTP) in sensory-evoked potentials. Journal of the Royal Society of New Zealand, 51(1), 24–40. https://doi.org/10.1080/03036758.2020.1780274

Kleeva, D. F., Rebreikina, A. B., Soghoyan, G. A., Kostanian, D. G., Neklyudova, A. N., & Sysoeva, O. V. (2022). Generalization of sustained neurophysiological effects of short-term auditory 13-Hz stimulation to neighbouring frequency representation in humans. European Journal of Neuroscience, 55(1), 175–188. https://doi.org/10.1111/ejn.15513

Klöppel, S., Lauer, E., Peter, J., Minkova, L., Nissen, C., Normann, C., Reis, J., Mainberger, F., Bach, M., & Lahr, J. (2015). LTP-like plasticity in the visual system and in the motor system appear related in young and healthy subjects [Original Research]. Frontiers in Human Neuroscience, 9. https://doi.org/10.3389/fnhum.2015.00506

Kompus, K., & Westerhausen, R. (2018). Increased MMN amplitude following passive perceptual learning with LTP-like rapid stimulation. Neuroscience Letters, 666, 28–31. https://doi.org/10.1016/j.neulet.2017.12.035

Lahr, J., Peter, J., Bach, M., Mader, I., Nissen, C., Normann, C., Kaller, C. P., & Klöppel, S. (2014). Heterogeneity of stimulus-specific response modification—an fMRI study on neuroplasticity [Original Research]. Frontiers in Human Neuroscience, 8. https://doi.org/10.3389/fnhum.2014.00695

Lei, G., Zhao, Z., Li, Y., Yu, L., Zhang, X., Yan, Y., Ma, X., Wang, Q., Wang, K., Zhang, D., Shen, W., Qiao, Y., & Yang, S. (2017). A method to induce human cortical long-term potentiation by acoustic stimulation. Acta Oto-Laryngologica, 137(10), 1069–1076. https://doi.org/10.1080/00016489.2017.1332428

Lengali, L., Hippe, J., Hatlestad-Hall, C., Rygvold, T. W., Sneve, M. H., & Andersson, S. (2021). Sensory-Induced Human LTP-Like Synaptic Plasticity – Using Visual Evoked Potentials to Explore the Relation Between LTP-Like Synaptic Plasticity and Visual Perceptual Learning [Original Research]. Frontiers in Human Neuroscience, 15. https://doi.org/10.3389/fnhum.2021.684573

Lømo, T. (1966). Frequency potentiation of excitatory synaptic activity in dentate area of hippocampal formation. Acta Physiologica Scandinavica, 68(suppl. 277), 128.

Lømo, T. (2003). The discovery of long-term potentiation. Philosophical Transactions of the Royal Society of London. Series B: Biological Sciences, 358, 617–620. doi: https://doi.org/10.1098/rstb.2002.1226

Lopez-Calderon, J., & Luck, S. J. (2014). ERPLAB: An open-source toolbox for the analysis of event-related potentials [Technology Report]. Frontiers in Human Neuroscience, 8(213). https://doi.org/10.3389/fnhum.2014.00213

McClaskey, C. M., Dias, J. W., Dubno, J. R., & Harris, K. C. (2018). Reliability of Measures of N1 Peak Amplitude of the Compound Action Potential in Younger and Older Adults. Journal of Speech, Language, and Hearing Research, 61(9), 2422–2430. https://doi.org/10.1044/2018_JSLHR-H-18-0097

McNair, N. A., Clapp, W. C., Hamm, J. P., Teyler, T. J., Corballis, M. C., & Kirk, I. J. (2006). Spatial frequency-specific potentiation of human visual-evoked potentials. NeuroReport, 17(7), 739–741. https://doi.org/10.1097/01.wnr.0000215775.53732.9f

Mears, R. P., & Spencer, K. M. (2012). Electrophysiological Assessment of Auditory Stimulus-Specific Plasticity in Schizophrenia. Biological Psychiatry, 71(6), 503–511. https://doi.org/10.1016/j.biopsych.2011.12.016

Moore, D., Ikuta, T., & Loprinzi, P. D. (2020). The Effects of Human Visual Sensory Stimuli on N1b Amplitude: An EEG Study. Journal of Clinical Medicine, 9(9), 2837. https://www.mdpi.com/2077-0383/9/9/2837

Moore, D., & Loprinzi, P. D. (2021). The association of self-reported physical activity on human sensory long-term potentiation. AIMS neuroscience, 8(3), 435–447. https://doi.org/10.3934/Neuroscience.2021023

Narne, V. k., & Vanaja, C. (2008). Speech identification and cortical potentials in individuals with auditory neuropathy [journal article]. Behavioral and Brain Functions, 4(1), 15. https://doi.org/10.1186/1744-9081-4-15

Normann, C., Schmitz, D., Fürmaier, A., Döing, C., & Bach, M. (2007). Long-Term Plasticity of Visually Evoked Potentials in Humans is Altered in Major Depression. Biological Psychiatry, 62(5), 373–380. https://doi.org/10.1016/j.biopsych.2006.10.006

Paternoster, R., Brame, R., Mazerolle, P., & Piquero, A. (1998). Using the correct statistical test for the equality of regression coefficients. Criminology, 36(4), 859–866. https://doi.org/10.1111/j.1745-9125.1998.tb01268.x

Peirce, J., Gray, J. R., Simpson, S., MacAskill, M., Höchenberger, R., Sogo, H., Kastman, E., & Lindeløv, J. K. (2019). PsychoPy2: Experiments in behavior made easy. Behavior Research Methods, 51(1), 195–203. https://doi.org/10.3758/s13428-018-01193-y

Rebreikina, A. B., Kleeva, D. F., Soghoyan, G. A., & Sysoeva, O. V. (2021). Effects of Auditory LTP-Like Stimulation on Auditory Stimulus Processing. Neuroscience and Behavioral Physiology, 51(9), 1323–1329. https://doi.org/10.1007/s11055-021-01197-w

Rosenthal, R. (1979). The file drawer problem and tolerance for null results. Psychological Bulletin, 86(3), 638–641. https://doi.org/10.1037/0033-2909.86.3.638

Rosenthal, R. (1991). Meta-analytic procedures for social research. Sage Publications.

Rosnow, R. L., Rosenthal, R., & Rubin, D. B. (2000). Contrasts and Correlations in Effect-Size Estimation. Psychological Science, 11(6), 446–453. https://doi.org/10.1111/1467-9280.00287

Ross, R. M., McNair, N. A., Fairhall, S. L., Clapp, W. C., Hamm, J. P., Teyler, T. J., & Kirk, I. J. (2008). Induction of orientation-specific LTP-like changes in human visual evoked potentials by rapid sensory stimulation. Brain Research Bulletin, 76(1), 97–101. https://doi.org/10.1016/j.brainresbull.2008.01.021

Rygvold, T. W., Hatlestad-Hall, C., Elvsåshagen, T., Moberget, T., & Andersson, S. (2020). Do visual and auditory stimulus-specific response modulation reflect different mechanisms of neocortical plasticity? [https://doi.org/10.1111/ejn.14964]. European Journal of Neuroscience, 53(4), 1072–1085. https://doi.org/10.1111/ejn.14964

Sanders, P. J., Thompson, B., Corballis, P. M., Maslin, M., & Searchfield, G. D. (2018). A review of plasticity induced by auditory and visual tetanic stimulation in humans. European Journal of Neuroscience, 48(4), 2084–2097. https://doi.org/10.1111/ejn.14080

Smallwood, N., Spriggs, M. J., Thompson, C. S., Wu, C. C., Hamm, J. P., Moreau, D., & Kirk, I. J. (2015). Influence of Physical Activity on Human Sensory Long-Term Potentiation. Journal of the International Neuropsychological Society, 21(10), 831–840. https://doi.org/10.1017/S1355617715001095

Spriggs, M. J., Cadwallader, C. J., Hamm, J. P., Tippett, L. J., & Kirk, I. J. (2017). Age-related alterations in human neocortical plasticity. Brain Research Bulletin, 130, 53–59. https://doi.org/10.1016/j.brainresbull.2016.12.015

Spriggs, M. J., Sumner, R. L., McMillan, R. L., Moran, R. J., Kirk, I. J., & Muthukumaraswamy, S. D. (2018). Indexing sensory plasticity: Evidence for distinct Predictive Coding and Hebbian learning mechanisms in the cerebral cortex. NeuroImage, 176, 290–300. https://doi.org/10.1016/j.neuroimage.2018.04.060

Spriggs, M. J., Thompson, C. S., Moreau, D., McNair, N. A., Wu, C. C., Lamb, Y. N., McKay, N. S., King, R. O. C., Antia, U., Shelling, A. N., Hamm, J. P., Teyler, T. J., Russell, B. R., Waldie, K. E., & Kirk, I. J. (2019). Human Sensory LTP Predicts Memory Performance and Is Modulated by the BDNF Val66Met Polymorphism [Original Research]. Frontiers in Human Neuroscience, 13. https://doi.org/10.3389/fnhum.2019.00022

Sumner, R. L., McMillan, R., Spriggs, M. J., Campbell, D., Malpas, G., Maxwell, E., Deng, C., Hay, J., Ponton, R., Kirk, I. J., Sundram, F., & Muthukumaraswamy, S. D. (2020). Ketamine Enhances Visual Sensory Evoked Potential Long-term Potentiation in Patients With Major Depressive Disorder. Biological Psychiatry: Cognitive Neuroscience and Neuroimaging, 5(1), 45–55. https://doi.org/10.1016/j.bpsc.2019.07.002

Sumner, R. L., Spriggs, M. J., McMillan, R. L., Sundram, F., Kirk, I. J., & Muthukumaraswamy, S. D. (2018). Neural plasticity is modified over the human menstrual cycle: Combined insight from sensory evoked potential LTP and repetition suppression. Neurobiology of Learning and Memory, 155, 422–434. https://doi.org/10.1016/j.nlm.2018.08.016

Sumner, R. L., Spriggs, M. J., Muthukumaraswamy, S. D., & Kirk, I. J. (2020). The role of Hebbian learning in human perception: a methodological and theoretical review of the human Visual Long-Term Potentiation paradigm. Neuroscience & Biobehavioral Reviews, 115, 220–237. https://doi.org/10.1016/j.neubiorev.2020.03.013

Suppa, A., Huang, Y. Z., Funke, K., Ridding, M. C., Cheeran, B., Di Lazzaro, V., Ziemann, U., & Rothwell, J. C. (2016). Ten Years of Theta Burst Stimulation in Humans: Established Knowledge, Unknowns and Prospects. Brain Stimulation, 9(3), 323–335. https://doi.org/10.1016/j.brs.2016.01.006

Team, R. C. (2022). R: A language and environment for statistical computing. In R Foundation for Statistical Computing. https://www.R-project.org/

Teyler, T. J., Hamm, J. P., Clapp, W. C., Johnson, B. W., Corballis, M. C., & Kirk, I. J. (2005). Long-term potentiation of human visual evoked responses. European Journal of Neuroscience, 21(7), 2045–2050. doi:https://doi.org/10.1111/j.1460-9568.2005.04007.x

Tremblay, K. L., Kraus, N., McGee, T., Ponton, C., & Otis, B. (2001). Central Auditory Plasticity: Changes in the N1-P2 Complex after Speech-Sound Training. Ear and Hearing, 22(2), 79–90. http://journals.lww.com/ear-hearing/Fulltext/2001/04000/Central_Auditory_PlasticityChanges_in_the_N1_P2.1.aspx

Valstad, M., Moberget, T., Roelfs, D., Slapø, N. B., Timpe, C. M. F., Beck, D., Richard, G., Sæther, L. S., Haatveit, B., Skaug, K. A., Nordvik, J. E., Hatlestad-Hall, C., Einevoll, G. T., Mäki-Marttunen, T., Westlye, L. T., Jönsson, E. G., Andreassen, O. A., & Elvsåshagen, T. (2020). Experience-dependent modulation of the visual evoked potential: Testing effect sizes, retention over time, and associations with age in 415 healthy individuals. NeuroImage, 223, 117302. https://doi.org/10.1016/j.neuroimage.2020.117302

Valstad, M., Roelfs, D., Slapø, N. B., Timpe, C. M. F., Rai, A., Matziorinis, A. M., Beck, D., Richard, G., Sæther, L. S., Haatveit, B., Nordvik, J. E., Hatlestad-Hall, C., Einevoll, G. T., Mäki-Marttunen, T., Haram, M., Ueland, T., Lagerberg, T. V., Steen, N. E., Melle, I., … Elvsåshagen, T. (2021). Evidence for Reduced Long-Term Potentiation-Like Visual Cortical Plasticity in Schizophrenia and Bipolar Disorder. Schizophrenia Bulletin, 47(6), 1751–1760. https://doi.org/10.1093/schbul/sbab049

Viechtbauer, W., & Cheung, M. W.-L. (2010). Outlier and influence diagnostics for meta-analysis. Research Synthesis Methods, 1(2), 112–125. https://doi.org/10.1002/jrsm.11

Wijtenburg, S. A., West, J., Korenic, S. A., Kuhney, F., Gaston, F. E., Chen, H., Roberts, M., Kochunov, P., Hong, L. E., & Rowland, L. M. (2017). Glutamatergic metabolites are associated with visual plasticity in humans. Neuroscience Letters, 644, 30–36. https://doi.org/10.1016/j.neulet.2017.02.020

Wilson, J. F., Lodhia, V., Courtney, D. P., Kirk, I. J., & Hamm, J. P. (2017). Evidence of hyper-plasticity in adults with Autism Spectrum Disorder. Research in Autism Spectrum Disorders, 43–44, 40-52. https://doi.org/10.1016/j.rasd.2017.09.005

Wischnewski, M., & Schutter, D. J. L. G. (2015). Efficacy and Time Course of Theta Burst Stimulation in Healthy Humans. Brain Stimulation, 8(4), 685–692. https://doi.org/10.1016/j.brs.2015.03.004

Wynn, J. K., Roach, B. J., McCleery, A., Marder, S. R., Mathalon, D. H., & Green, M. F. (2019). Evaluating visual neuroplasticity with EEG in schizophrenia outpatients. Schizophrenia Research, 212, 40–46. https://doi.org/10.1016/j.schres.2019.08.015

Zaehle, T., Clapp, W. C., Hamm, J. P., Meyer, M., & Kirk, I. J. (2007). Induction of LTP-like changes in human auditory cortex by rapid auditory stimulation: an FMRI study. Restorative Neurology and Neuroscience, 25(3-4), 251–259.

Zak, N., Moberget, T., Bøen, E., Boye, B., Waage, T. R., Dietrichs, E., Harkestad, N., Malt, U. F., Westlye, L. T., Andreassen, O. A., Andersson, S., & Elvsåshagen, T. (2018). Longitudinal and cross-sectional investigations of long-term potentiation-like cortical plasticity in bipolar disorder type II and healthy individuals. Translational Psychiatry, 8(1), 103. https://doi.org/10.1038/s41398-018-0151-5

